# An early Cambrian polyp reveals an anemone-like ancestor for medusozoan cnidarians

**DOI:** 10.1101/2021.12.24.474121

**Authors:** Yang Zhao, Luke A. Parry, Jakob Vinther, Frances S. Dunn, Yu-jing Li, Fan Wei, Xian-guang Hou, Pei-yun Cong

## Abstract

Extant cnidarians are a disparate phylum of non-bilaterians and their diploblastic body plan represents a key step in animal evolution. Anthozoans (anemones, corals) are benthic polyps, while adult medusozoans (jellyfishes) are dominantly pelagic medusae. A sessile polyp is present in both groups and is widely conceived as the ancestral form of their last common ancestor. However, the nature and anatomy of this ancestral polyp, particularly of medusozoans, are controversial, owing to the divergent body plans of both groups in the extant lineages and the rarity of medusozoan soft tissues in the fossil record. Here we redescribe the enigmatic *Conicula striata* Luo et Hu from the early Cambrian Chengjiang biota, south China, which has previously been interpreted as a polyp, lophophorate or deuterostome. We show that *C. striata* possessed features of both anthozoans and medusozoans. Its stalked polyp and fully encasing conical, annulated organic skeleton (periderm) are features of medusozoans. However, the gut is partitioned by ∼28 mesenteries, and has a tubular pharynx, resembling anthozoans. Our phylogenetic analysis recovers *C. striata* as a stem medusozoan, indicating that the enormously diverse medusozoans were derived from an anemone-like ancestor, with the pharynx lost and number of mesenteries reduced prior to the origin of crown group Medusozoa.

## Introduction

Among non-bilaterian animals, Cnidaria is the phylum with the most species richness as well as the most disparate range of morphologies (Daly et al., 2007). Members of Cnidaria share the presence of tentacles with stinging cells (cnidocytes) used in prey capture, a blind gastric cavity that is often partitioned by mesenteries/septa and a sessile polypoid phase in at least part of their lifecycle in most groups (Hyman, 1940). The phylum mainly includes two monophyletic subgroups: Anthozoa and Medusozoa (Marques and Collins, 2004; Schuchert, 1993). Anthozoans are primarily benthic, polypoid animals encompassing widely known groups such as sea anemones and corals (Brusca et al., 2016), whereas medusozoans usually have a biphasic lifestyle, with sessile polyps giving rise to swimming medusae (jellyfishes) via asexual reproduction (Collins, 2002). Interpretations of cnidarian phylogeny support a scenario in which the common ancestor of the crown group was a sessile polyp, and the swimming medusa represents a synapomorphy of Medusozoa (Collins et al., 2006; Kayal et al., 2018; Marques and Collins, 2004; McFadden et al., 2021). However, the anatomy of the polyps diverges in living anthozoans and medusozoans in several respects (Daly et al., 2007; Technau and Steele, 2011), rendering the inference of their ancestral forms uncertain.

Anthozoan polyps possess a tubular pharynx (actinopharynx) that extends from the mouth to a gastric cavity that is partitioned by well-developed mesenteries (Daly et al., 2007). The pharynx contains either one or two ciliated siphonoglyphs that impart a bilateral/biradial symmetry (Malakhov, 2016). In contrast, medusozoan polyps are relatively small and possess an exoskeleton called periderm, most of which are made of chitin (Mendoza-Becerril et al., 2016). This organic skeleton can encase the entire body of the polyp in some lineages such as coronate Scyphozoa, or be reduced to just the basal portion of the polyp (Mendoza-Becerril et al., 2016). In medusozoan polyps, the mouth is usually extended away from the body on a protuberance referred to as the scyphopharynx or hypostome in different subgroups. The gastric cavity is not always portioned by septa, as they are absent in the main subclades of hydropolyps (Bouillon and Boero, 2000) and cubopolyps (Chapman, 1978).

Molecular clock estimates suggest that the cnidarian crown groups radiated in the Ediacaran or the Cryogenian (dos Reis et al., 2015; Park et al., 2012), but cnidarian fossils before the Cambrian are rare and/or controversial (Liu et al., 2014; Van Iten et al., 2014; Van Iten et al., 2016). Cambrian deposits, however, yield a wealth of cnidarian fossils that exceptionally preserved delicate soft tissues (Cartwright et al., 2007; Conway Morris, 1993; Han et al., 2016; Hou et al., 2005) and even their early developmental stages (Dong et al., 2016), and are therefore crucial to understand the origin and early evolution of cnidarian clades. Here, we redescribe the enigmatic *Conicula striata* Luo et Hu (Luo et al., 1999) from the Cambrian (Epoch 2, Age 3) Chengjiang biota from Yunnan Province, south China. *C. striata* has previously been interpreted as a lophophorate (spiralian) (Luo et al., 1999), an actinarian (cnidarian) (Hu, 2005) or a phlogitid (presumed deuterostome) (Caron et al., 2010). However, with only one specimen reported, *C. striata* has remained as one of the most poorly understood early Cambrian fossils. In this study, we depict the detailed morphology of *C. striata* based 51 exquisitely preserved specimens, revealing mosaic morphological characteristics as seen in both extant anthozoans and medusozoans. These features bring *C. striata* in the stem of Medusozoa, unveiling the earliest medusozoan polyp as an anemone-like form encased in an extensive periderm.

## Results

### Systematic Palaeontology

> Phylum Cnidaria Verrill, 1865

> Stem group Medusozoa Petersen, 1979

> *Conicula striata* Luo et Hu, 1999

> (Figures 1-3, S1-S7)

### Occurrence

*Conicula striata* occurs in Cambrian Series 2, Stage 3, the local lithostratigraphic unit is the Yu’anshan Member of the Chiungchussu Formation, corresponding to the *Eoredlichia-Wutingaspis* trilobite biozone (Babcock and Zhang, 2001).

**Amended diagnosis** (amended from *Luo et al., 1999*, p. 87). Solitary polypoid cnidarian encased in a conical, annulated periderm. The polyp possesses unbranched, flexible circumoral tentacles that can protrude outwards from the distal end of the periderm, and a blind digestive tract consisting of an elongate, tubular pharynx and a mesentery-partitioned gastric cavity.

### Description

Specimens of *Conicula striata* usually exhibit a conical shape in lateral view (Figures 1, S1, and S2) and occasionally a circular cross section in oblique-lateral view (Figure S3). The body is 10-38 mm long with maximum width ranging between 4-17 mm (width/length ratio among complete specimens is 0.38-0.45). Some specimens preserve evidence of both a rigid, external skeleton (periderm) (Figure 2), circumoral tentacles (Figure 3A-C) and a columnar body with internal anatomical features (Figure 3D-E, G-H, J-L).

**Figure 1.**
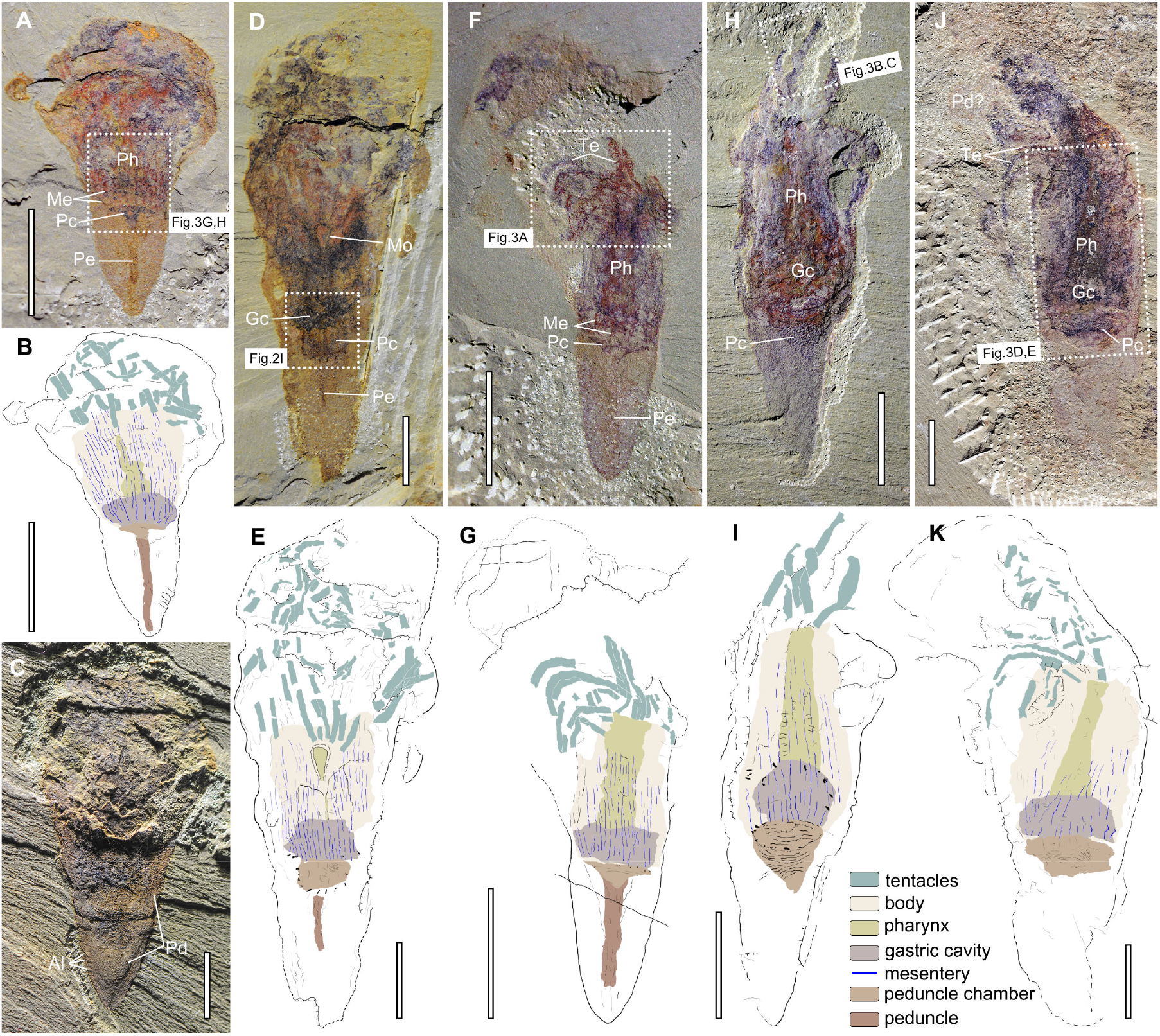
General morphology of *C. striata*. (A-B) YKLP 13212a. (C) YKLP 13213a. (D-E) YKLP 13215a. (F-G) YKLP 13288a. (H-I) YKLP 13484a. (J-K) YKLP 13485a. Al, annulation; Gc, gastric cavity; Me, mesentery; Mo, mouth; Pc, peduncle chamber; Pd, periderm; Pe, peduncle; Ph, pharynx; Te, tentacle. Scale bars: 5 mm (A-K). See also Figures S1-S7.

**Figure 2.**
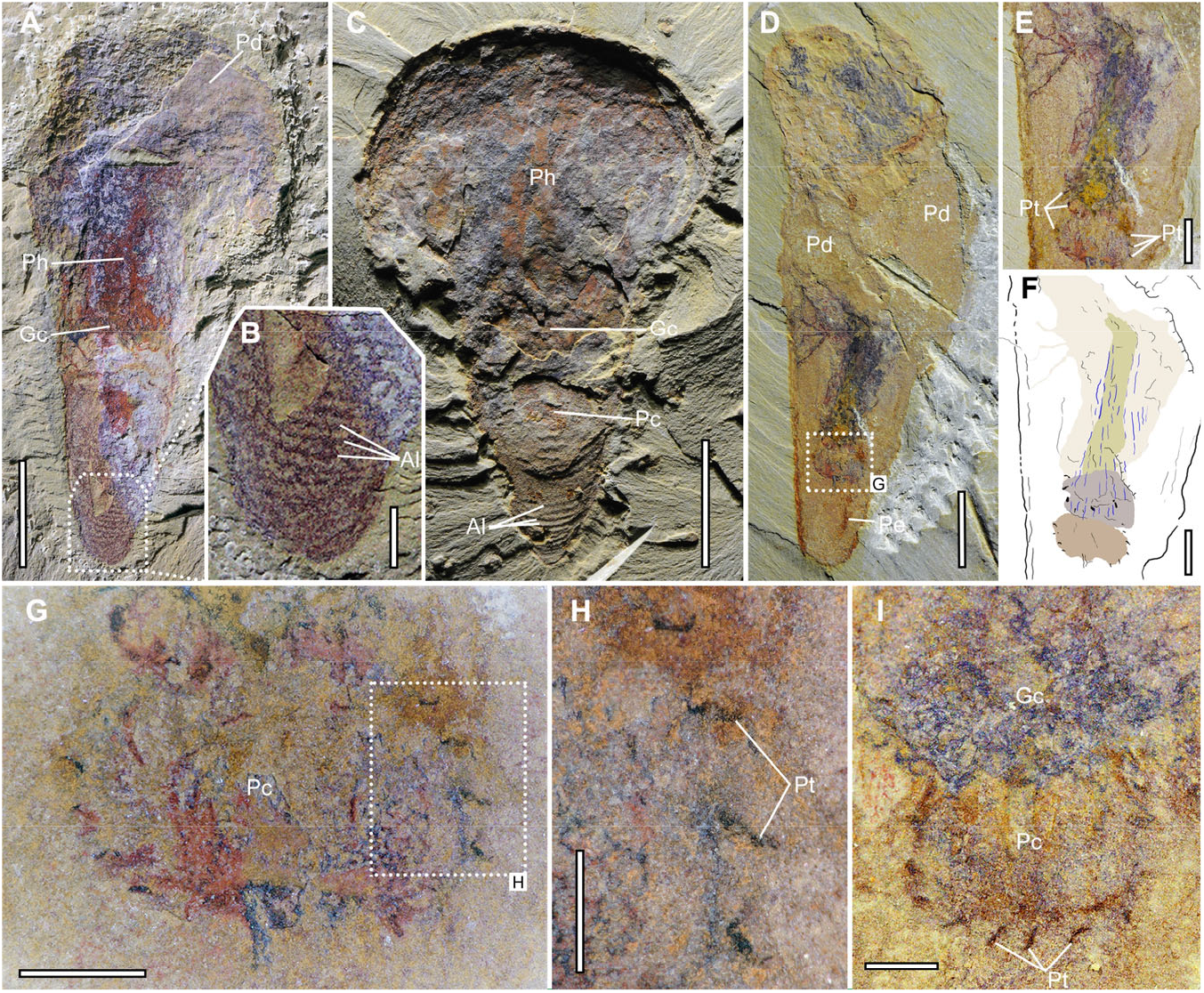
Exoskeleton/periderm of *C. striata*. (A) YKLP 13488, complete specimen showing smooth periderm in the broken globular region. (B) Magnification of the apical portion of the skeleton, showing parallel annulations. (C) YKLP 13214b, under low angle light, showing slight relief of annulations in the proximal portion. (D-H) YKLP 13210a, showing polyp soft tissues within the periderm. (E-F) Close-up of the soft parts of the polyp and interpretative drawing (colour scheme as in Figure 1). (G-H) Close-up of the peduncle chamber, encircled by dark, spine-like peridermal teeth. (I) YKLP 13215a, showing the gastric cavity and peridermal teeth. Al, annulation; Gc, gastric cavity; Pc, peduncle chamber; Pd, periderm; Pe, peduncle; Ph, pharynx; Pt, peridermal teeth. Scale bars: 500 μm (H); 1 mm (B, G, I); 2 mm (E, F); 5 mm (A, C, D). See also Figure S2.

**Figure 3.**
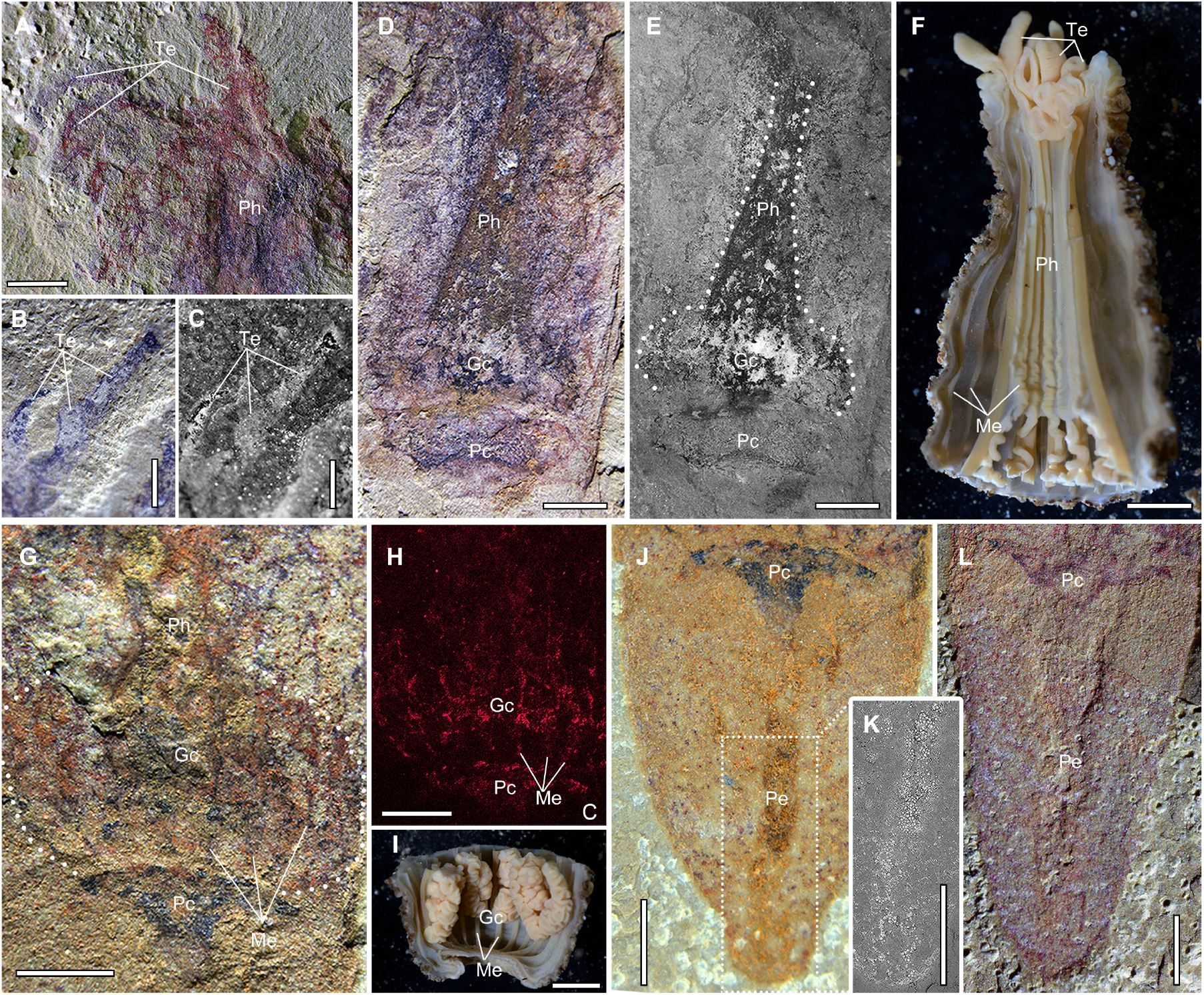
Internal anatomy of *C. striata*. (A) YKLP 13288a, showing circumoral tentacles. (B-C) YKLP 13484a, close-up of tentacles, showing flexible, unbranched features, in direct light (B) and fluorescent light (C). (D-E) YKLP 13485a, details of internal structures, showing a tubular pharynx, gastric cavity and peduncle chamber. (F) Extant sea anemone *Cactosoma abyssorum*(Sanamyan et al., 2016), longitudinal section of the distal half of the body, showing circumoral tentacles, an actinopharynx and mesenteries.(G-H) YKLP 13212a, details of the gastric cavity partitioned by longitudinal, dark lines (mesenteries), imaged in direct light (G) and elemental map of carbon (H). (I) Living sea anemone *Cactosoma abyssorum*(Sanamyan et al., 2016), longitudinal section of the proximal part of the body, showing a gastric cavity partitioned by more than eight mesenteries. (J) YKLP 13212a, close-up of the lower conical portion, showing a tube-like peduncle extended from the dark peduncle chamber. (K) Magnification of the peduncle using backscatter SEM, showing abundant weathered pyrite granules gathered in the peduncle. (L) YKLP 13288a, close-up of the lower conical portion, showing slight relief of the peduncle. Gc, gastric cavity; Me, mesentery; Pc, peduncle chamber; Pe, peduncle; Ph, pharynx; Te, tentacle. Scale bars: 1 mm (A-C, G, H, J-L); 2 mm (D-F, I). See also Figures S4 and S5.

The external skeleton appears to have been originally inflexible and robust, and fully encloses the aboral portion of the body (“Pd”, Figures 2A, C, D and S2D-F). The skeleton forms a conical structure in the proximal portion, which during growth expanded and then narrowed to form a globular chamber, resulting in a shape resembling a classic, well-loaded ice cream cone. It is ornamented by parallel annulations (“Al”, Figure 2A-C), with slight relief under low angle light (Figures 2C and S1D) which are most visible in the aboral section. The space between adjacent annulations is around 0.2-0.4 mm. The appearance of the exoskeleton is consistent with the periderm of medusozoan cnidarians. Dark, spine-like structures project into the interior of the periderm. They are arranged in one whorl and surround internal features, particularly the region housing the digestive tract, and are termed here as peridermal teeth (“Pt”, Figures 2D-I, S2A, and S6E-G).

Soft tentacles are exhibited at the distal part of the body (“Te”, Figures 1F, H, J, 3A-C, and S1E, F), either approximately straight (e.g. Figure 1H) or curved (e.g. Figure 1F, J) and vary in aspect ratio, suggesting that they were capable of protruding and retracting. The tentacles are smooth, with no identifiable branches or pinnules (Figures 3A-C and S1E, F). When present, the tentacles are buckled and intertwined as a dark organic-rich mass within the globular chamber, making it difficult to trace each tentacle (Figures 1A-E, S2C, E, F, S5B, and S6A-D). It is challenging to surmise the total number of tentacles due to twisting and/or superposition of multiple tentacles as well as incomplete exposure and varying degrees of decay.

The tentacle base surrounds a disc with a central dark, tongue-shaped structure, likely representing the mouth opening (“Mo”, Figures 1D, E and S5C, D). An elongate, narrow tubular structure (about 6.5 mm and 7.7 mm long in YKLP 13484a and 13485a, respectively) connects the opening to the lower part of the body (“Ph”, Figures 1F-K, 3D, E, and S5E-H), in which an oval-shaped structure with a dark outline expands to near the body margin (“Gc”, Figures 1, 3D, E, and S5G, H). The tapering band and oval-shaped structure are distinct from other regions by their preservation in a red-brown or dark colour when viewed with cross-polarised light or fluorescence microscopy, respectively (Figures 3D, E, S3A, D, S4D, E, and S5G, H), and their higher amounts of carbon and iron in elemental maps (Figures S4F, G, S6E, G, and S7F, G). The tubular structure and oval-shaped structure are interpreted as a pharynx and a gastric cavity, respectively, based on their shape, size and position within the animal body (Figure 3F, I).

The gastric cavity varies in size and shape in different specimens, but frequently preserves evidence that was partitioned by dark, longitudinal lines (“Me”, Figures 3G, H, S2C, S3A-C, S4A-F, and S6A-D). In YKLP 13212a, these dark lines are nearly parallel with each other and extend from the base of the oval-shaped structure to the distal region of the columnar body (Figures 3G, H and S4D-F). The spacing between two adjacent lines in the central region is ∼0.39 mm, suggesting a total of 28 mesenteries or so (calculated by dividing this interval by the body circumference≈11.3 mm). However, dark lines are sometimes inconspicuous, with some superposition from either side of the body, making their precise number difficult to determine across specimens. These dark lines are carbonaceous in preservation (Figure 3H) and interpreted as mesenteries/gastric septa according to their appearance and position (Figure 3I).

A further dark patch, with a circular, crescent or triangular outline, attaches at the base of the gastric cavity (“Pc”, Figures 1 and S2). It contains a high abundance of carbon as revealed by elemental maps (Figure S6H, I). A slightly curved, narrower ribbon-like structure (“Pe”, Figures 1A, D, F, 3J, L, and S4A-C), sometimes with modest relief (Figure 3L), protrudes from the bottom of the dark patch to the proximal end of the periderm (e.g. YKLP 13212 and 13288). In YKLP 13212, it is roughly parallel sided (∼0.34 mm wide), and contains abundant weathered pyrite granules (Figure 3K). The dark patch and ribbon-like structure are herein interpreted as a peduncle chamber and a peduncle, respectively.

### Phylogenetic position

Our phylogenetic analysis incorporating *Conicula striata* includes 99 taxa and 304 characters scored for a diversity of living and fossil cnidarians and outgroups. Analysis of this dataset without any topological constraints recovers *C. striata* in the medusozoan stem group and all other tubular fossils in a clade that is in a polytomy with extant medusozoan taxa (Figures 4C and S8A). When removing these tubular fossils, *C. striata* is still recovered in the medusozoan stem group while extant medusozoan taxa are monophyletic (Figure S8B). This analysis also recovers the paraphyly of Scyphozoa, a result that is not found in recent molecular phylogenies but has been found in many recent morphological phylogenies(Duan et al., 2017; Zhao et al., 2019). Constraining the in-group relationships of cnidarians still recovers *C. striata* in the stem group of Medusozoa (Figure S8C).

**Figure 4.**
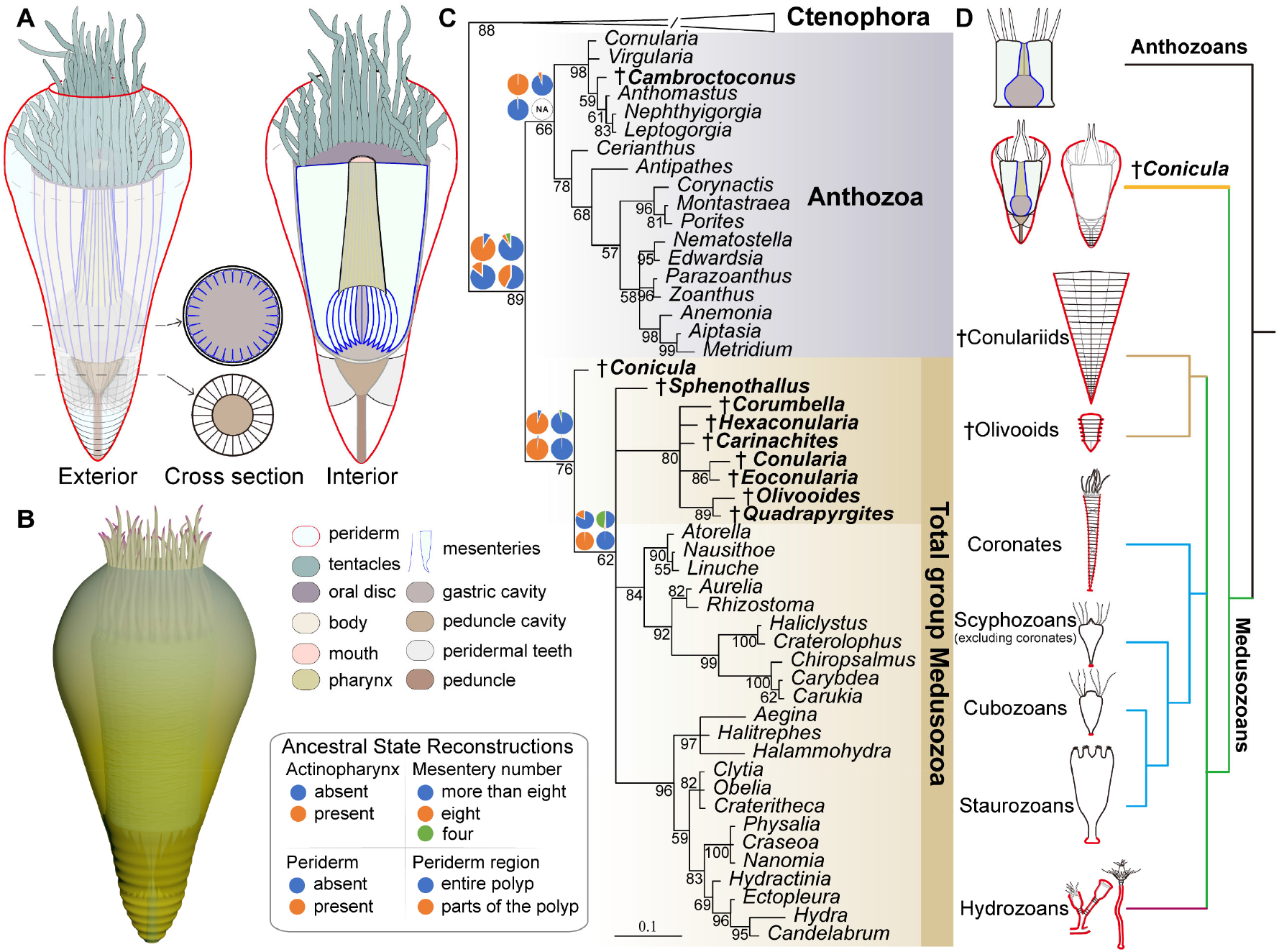
Reconstruction and phylogenetic analysis. (A) Technical reconstruction showing the exterior gross morphology, interior anatomy and cross section of gastric cavity and peduncle chamber. The colour scheme is described to the top right of (B). (B) Three-dimensional model of *C. striata.* (C) Bayesian phylogenetic analysis (304 characters, 99 taxa, mkv+gamma model), *C. striata* is resolved as a stem group medusozoan. The fossil taxa are indicated by dagger symbol. Pie charts illustrate ancestral states from Bayesian analysis. Numbers at the nodes are posterior probabilities, and the scale bar is the expected number of substitutions per site. (D) Simplified cladogram of cnidarians, showing that *C. striata* has mosaic characters of anthozoan (a high number of mesenteries and a tubular pharynx) and medusozoan (an annulated periderm and teeth) polyps. See also Figure S8 for full results and additional information.

## Discussion

### *C. striata* is a stem medusozoan with a mixed anthozoan-medusozoan body plan

*Conicula striata* was erected in 1999 based on one incomplete specimen preserving a tentacular, annulated conical body (Luo et al., 1999). It has since been variously interpreted as a lophophorate (Luo et al., 1999), a sea anemone (Hu, 2005) or a deuterostome (Caron et al., 2010). Our 51 new specimens corroborate the validity of this genus and further provide new insights into its morphology, phylogenetic position and evolutionary significance. *C. striata* possesses an expansive, partitioned gastric cavity with a single opening (Figure 3D-E, G-H), unbranched, circumoral tentacles (Figure 1F, H, J) and a radially symmetrical body (Figure S3). These characters are incompatible with the previous interpretations as either a lophophorate or a deuterostome. Lophophorates possess a U-shaped gut (with a mouth and an anus) that is often conspicuously preserved in Chengjiang fossils (Zhang et al., 2013). Although blind guts are present in articulate brachiopods, *C. striata* shares no derived characters with brachiopods in any aspect of body construction. Cambroernids (presumed deuterostomes), a fossil group including *Herpetogaster*, phlogitids and eldoniids (Caron et al., 2010), also differ markedly as they have bilaterally arranged, paired, branched, even dendritic tentacles and a discrete, through gut (Caron et al., 2010; Hou et al., 2006).

The regular, longitudinal dark lines preserved in association to the columnar body wall and the gastric cavity (Figure 3G, H) are interpreted as mesenteries. Similar structures have been identified in Cambrian exceptionally preserved fossils before, such as *Archisaccophyllia* (Hou et al., 2005) and some dinomischiids (Zhao et al., 2019). Partitioning of the gastric cavity by mesenteries/septa is a widespread character in extant cnidarian polyps (Figure 3I) (Daly et al., 2007), with a higher number (at least 8) of well-developed mesenteries occurring in anthozoans while only 4 present in scyphopolyps and stauropolyps. In hydropolyps (Bouillon and Boero, 2000) and cubopolyps (Chapman, 1978) gastric septa are absent. The number of mesenteries observed in *C. striata* (∼28) is consistent with that in anthozoans. The elongate tubular structure of *C. striata* that extends from the gastric cavity to the oral disc and mouth corresponds in size and topological position to the actinopharynx of anthozoans (Figure 3D-F) (Daly et al., 2003; Daly et al., 2007), but is different from the scyphopharynx/hypostome of medusozoans which is an oral extension at the distal end of the body. The digestive system of *C. striata* therefore closely resembles the condition seen in extant anthozoans.

The exoskeleton preserves regularly arranged annulations in the proximal portion in some specimens (Figures 2A-C and S1A-D). While compaction could have enhanced their relief, they are unlikely to be artefactual due to the common occurrence across specimens. Such annulations resemble the growth lines caused by the marginal accretion of an exoskeleton, which are widely present in extant cnidarians (e.g. medusozoan periderm) and spiralians (such as molluscs, brachiopods and tubular polychaetes). Given that the polypoid body and internal structures of *C. striata* deviate from the body plans of bilaterians, we limit comparisons of the skeleton to those of cnidarians. Anthozoans also produce skeletons, which are found in living antipatharians, ceriantharians, scleractinians and octocorallians, and extinct rugose and tabulate corals, but presumably have multiple independent origins. Among these, *C. striata* only superficially resembles the ceriantharians, which produce tubes made of mucus and ptychocysts (Stampar et al., 2015). The accretionary exoskeleton of *C. striata* is directly comparable to the periderm of medusozoans and we infer that these features are homologous.

The dark, spine-like structures (Figure 2D-I) that encircle the basal portion of the columnar body are interpreted as the remains of peridermal teeth. Extant coronate polyps possess well-developed whorls of complex peridermal teeth, which are protrusions of the inner layer of the periderm towards the central chamber, to anchor the polypoid body (Jarms, 1991). Similar peridermal teeth also appear in tubular fossils as sheet-like ridges (cusps) in *Sphenothallus* (Dzik et al., 2017) or paired projections in *Olivooides* (Dong et al., 2016). These peridermal teeth are often repeated along the length of the tube wall (Jarms, 1991), but in *C. striata* they occur in a single whorl towards the aboral end of the tube, with the remainder of the lumen of the skeleton appearing smooth. In addition, the interpreted peduncle might also function for anchoring the polypoid body inside the theca, along with the peridermal teeth. In colonial hydropolyps, peduncle-like structures connect individual polyps together (Cartwright, 2004) and are therefore not readily comparable with that seen in *C. striata*.

*C. striata* is inferred here to have been a solitary organism with an annulated periderm fully encasing the polyp, evidencing a benthic and sessile lifestyle commonly seen among coronate and hydrozoan polyps. However, all specimens lack a holdfast at the aboral end and instead appear to taper naturally, with no evidence of attachment to other organisms or substrates, suggesting *C. striata* may have embedded the apex into the seafloor for anchoring, similar to some conulariids (Van Iten et al., 2013). Alternatively, *C. striata* may have been recumbent, but the conical skeleton does not curve to facilitate such a mode of life like that of horn corals (Scrutton, 1998). An alternative less plausible scenario is that *C. striata* was planktonic, with buoyancy provided by the inflated distal chamber, possibly having an intermediate lifestyle between the benthic polyps and pelagic medusae.

### Evolutionary significance

Historically, many tubular fossils from the Ediacaran-Cambrian have been interpreted as cnidarian polyps (Table S1)(Van Iten et al., 2014), such as the microfossils Olivooidae, Carinachitidae and Hexangulaconulariidae(Guo et al., 2020) as well as macrofossils *Corumbella* and conulariids(Van Iten et al., 2016). The features revealed in these tubular fossils, such as radial symmetry, transverse ribs/crests, peridermal teeth and a single opening, are the primary lines of evidence for a cnidarian interpretation, with particularly close comparisons made to the peridermal tubes of medusozoan polyps(Conway Morris and Chen, 1992; Dong et al., 2016; Van Iten et al., 2006; Zhu et al., 2000). Soft tissues are extremely scarce among these fossil tubes, and their cnidarian affinities and interpretations are accordingly not without previous controversy(Steiner et al., 2014; Walde et al., 2019). They are recovered as a paraphyletic grade of total group medusozoans in our Bayesian analyses (Figures 4C, S8). *C. striata* not only shares with those tubular fossils similar exoskeletal features, but also provides unique new evidence for the soft tissues of early medusozoans, such as mesenteries, the digestive tract and tentacles, characters that are not available from the overwhelming majority of tubular fossil taxa, shedding light on character state changes that occurred early in medusozoan evolutionary history.

*C. striata* shows a tubular pharynx, which is similar to the actinopharynx of anthozoans in topological location, inferred function and architecture (Figure 3D-F). The presence of an anthozoan-like pharynx in the medusozoan stem group is also supported in our ancestral state reconstruction (Figure 4C). It is recovered as a plesiomorphic trait of cnidarians (Figure 4C), indicating that the tubular pharynx in the stem group of Medusozoa is homologous with the actinopharynx in anthozoans, but it was subsequently lost prior to the origin of crown group medusozoans. Whether other tubular fossil taxa have an anthozoan-like pharynx is not known, which would depend on further findings of soft tissues in these groups. Moreover, *C. striata* possesses about twenty-eight mesenteries lining the gastric cavity, a feature commonly seen in extant hexacorallians, which is also recovered as being plesiomorphic for medusozoans and cnidarians in our analysis (Figure 4C).

We infer that the periderm is a true medusozoan synapomorphy(Mendoza-Becerril et al., 2016), but is absent in the common ancestor of Anthozoa, and probably of Cnidaria (Figure 4C). This inference is in congruence with earlier cnidarian fossils from the Ediacaran-Cambrian, in which all potential medusozoan polyps share a well-developed, annulated exoskeleton (periderm). In contrast, all known anthozoan fossil taxa lack a comparable exoskeleton/periderm (Han et al., 2010; Hou et al., 2005). A polyp encased fully by a periderm is recovered as a plesiomorphic trait of the medusozoan total group (Figure 4C) and this is only retained in living coronates and some members of Hydrozoa.

In light of this, we reconstruct the ancestral medusozoan as an anemone-like polyp, which possessed a tubular pharynx (actinopharynx) connecting the mouth and gastrovascular cavity that is partitioned by ∼28 mesenteries and unbranched tentacles, with the body encased fully by an annulated exoskeleton (periderm) (Figure 4A, B). The body plan of *C. striata* bridges the long-known morphological gap between living anthozoan and medusozoan polyps (Figure 4D), suggesting that several features previously regarded as anthozoan apomorphies (e.g. an actinopharyx) might have a deeper origin and are shared by stem medusozoans.

Given the anatomical simplicity in crown medusozoan polyps(Ruppert et al., 2004), we infer that several characters, including well-developed mesenteries and a tubular pharynx, experienced subsequent reduction and even total loss in some or all extant medusozoan lineages (Figure 4D). Only a few living medusozoan polyps (e.g. coronate scyphozoans) have a well-developed periderm and it is completely restricted to the lower body or podocyst in some lineages (e.g. staurozoans)(Mendoza-Becerril et al., 2016). Our analyses indicate that this is also the result of secondary reduction and a component of a broader trend of polyp simplification in Medusozoa where the lifecycle is now dominated by the medusa stage.

## Materials and Methods

### Materials

51 specimens were collected in fieldwork from 2014 to 2019 in Haikou area, Kunming, eastern Yunnan province, south China. They were checked and prepared under a Leica M205C stereomicroscope. A fine needle was used to remove the matrix and expose the fossils. The specimen size was measured with ImageJ 1.51j8. All specimens are housed in the Yunnan Key Laboratory for Palaeobiology (YKLP), Yunnan University, China. The holotype (He-f-6-5-112/113) is deposited in the Yunnan Institute of Geological Survey, Kunming, China.

### Photography

16 out of 51 specimens were figured. Photographs were taken either by a Canon EOS 5DS R digital camera mounted with Canon MP-E 100mm or 65mm macro lens (1-5X), using high/low angle cross-polarised light, or by a Leica DFC 5000 camera mounted on Leica M205C microscope (to obtain morphological details). Interpretative drawings were combined with camera lucida drawings done under Nikon SMZ1000 stereomicroscope and digital photographs. Fluorescence images were obtained using a Leica DFC7000 T digital camera linked to a Leica M205 FA fluorescence microscope. All figures were processed in Adobe Photoshop CC 2019 to adjust the levels, brightness and contrast. The reconstruction of the body plan was drafted in Adobe Illustrator CC 2019.

### Scanning electron microscopy (SEM)

SEM images were collected by a FEI Quanta 650 FEG in low-vacuum mode using an accelerating voltage of 15kV (30kV in Figure S7F). Elemental composition analyses were carried out with an EDAX Pegasus using accelerating voltages of 15kV (10kV in Figure S6I). All above analyses were performed in the Institute of Palaeontology, Yunnan University.

### Phylogenetic analysis

Phylogenetic analyses were performed in MrBayes 3.2.7 (Ronquist et al., 2012) under the mkv + gamma model (Lewis, 2001). 20,000,000 generations were requested, with analyses stopping automatically once the average deviation of split frequencies was <0.01. Ancestral character states for selected nodes were reconstructed in separate analyses using monophyly constraints also performed in MrBayes 3.2.7. Posterior probabilities for the character states at these nodes are plotted as pie charts at nodes shown in Figure 4C.

## Supplementary information

Supplementary information includes eight figures, one table and phylogenetic information (character descriptions and codes).

## Acknowledgments

We thank Qiang Ou (China University of Geosciences, Beijing) and Shixue Hu (Chengdu Center, China Geological Survey) for their insightful comments and suggestions, and Karen E. Sanamyan (Kamchatka Branch of Pacific Geographical Institute) for sharing extant sea anemones images in Figure 3F, I, along with Mengying Yin for creating the 3D model in Figure 4B, and Shangnan Zhang for her help with SEM analyses. This work was supported by the National Natural Science Foundation of China (42072019 and 42062001 to P.-Y.C. and F.W.), the Strategic Priority Research Program of Chinese Academy of Sciences (XDB26000000 to P.-Y.C. and Y.-J.L.) and State Key Laboratory of Palaeobiology and Stratigraphy (Nanjing Institute of Geology and Palaeontology, CAS) (193128 and 213111 to F.W. and Y.-J.L.). Y.Z. is supported by a graduate grant from China Scholarship Council (201907030012). F.S.D. is supported by fellowships from Merton College and 1851. L.A.P. is supported by an early career fellowship from St Edmund Hall.

## Author contributions

Y.Z. and P.-Y.C. designed research; Y.Z., P.-Y.C., X.-G.H., Y.-J.L. and F.W. collected fossil material; Y.Z., L.A.P. and J.V. performed research, analysed data and prepared all figures; Y.Z., F.S.D and L.A.P collated the morphological data and conducted phylogenetic analyses; Y.Z. and L.A.P. wrote the initial manuscript with significant input from J.V., P.-Y.C. and all other co-authors.

## Declaration of interests

The authors declare no competing interests.

## Supplementary Information

### This PDF file includes

Figures S1 to S8

Tables S1

Phylogenetic information

SI References

### Other supplementary materials for this manuscript include the following

Datasets S1 *Conicula* phylogenetic code

**Fig. S1.**
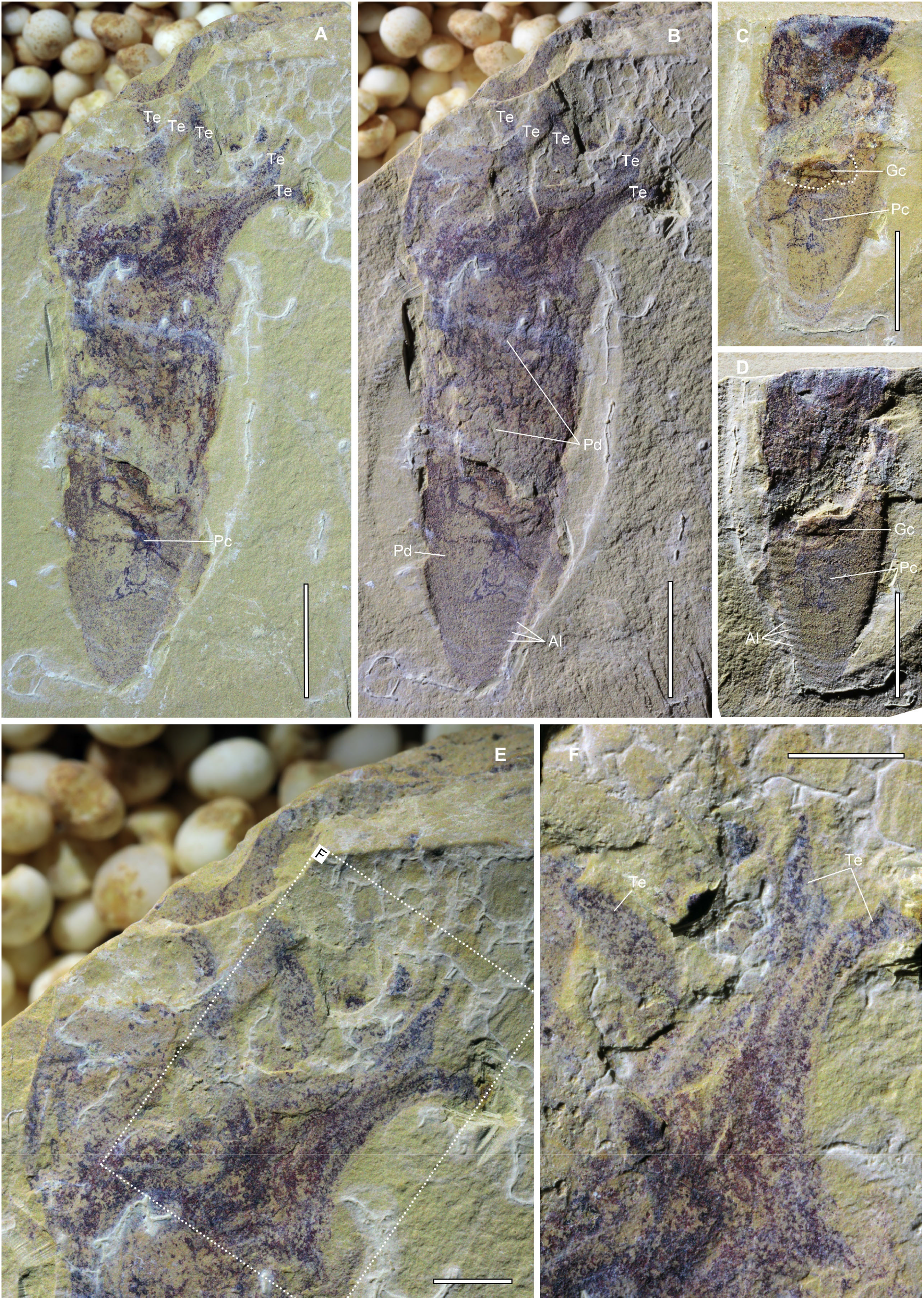
The holotype of *C. striata*. (A-B) The part of holotype, He-f-6-5-112. (C-D) The counterpart of holotype, He-f-6-5-113. Imaged under high angle (A, C) and low angle (B, D) direct light. (E) Magnification of the distal region, showing scattered tentacles. (F) Details of the unbranched tentacles. Al, annulation; Gc, gastric cavity; Pc, peduncle chamber; Pd, periderm; Te, tentacle. Scale bars: 2 mm (E and F); 5 mm (A-D).

**Fig. S2.**
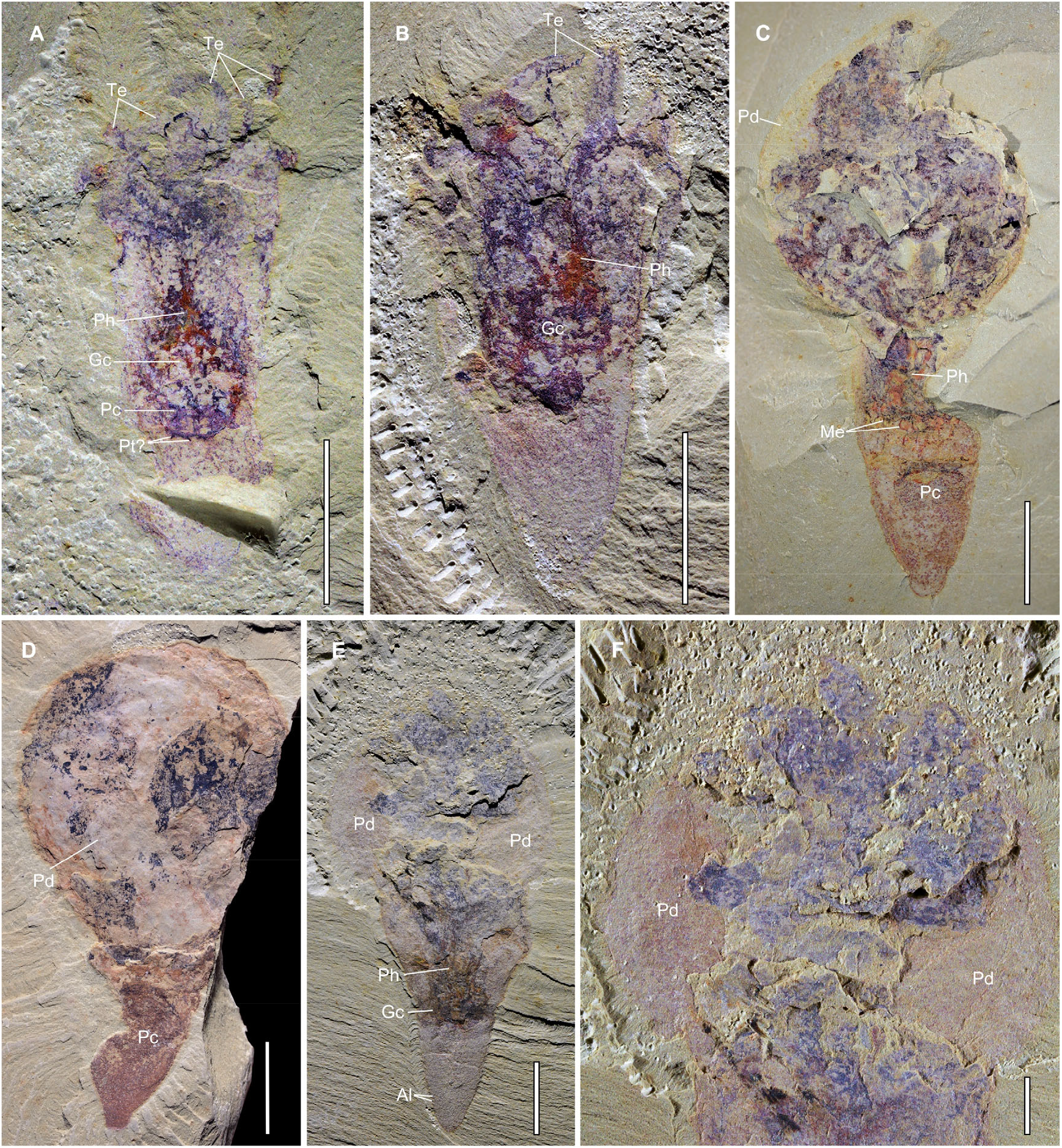
Additional specimens of *C. striata.* (A) YKLP 13486. (B) YKLP 13290. (C) YKLP 13494a. (D) YKLP 13490. (E) YKLP 13220; (F) Close-up of the globular region in E, the external periderm is exposed due to its broken preservation. Al, annulation; Gc, gastric cavity; Me, mesentery; Pc, peduncle chamber; Pd, periderm; Ph, pharynx; Pt, peridermal teeth; Te, tentacle. Scale bars: 2 mm (F); 5 mm (A-E).

**Fig. S3.**
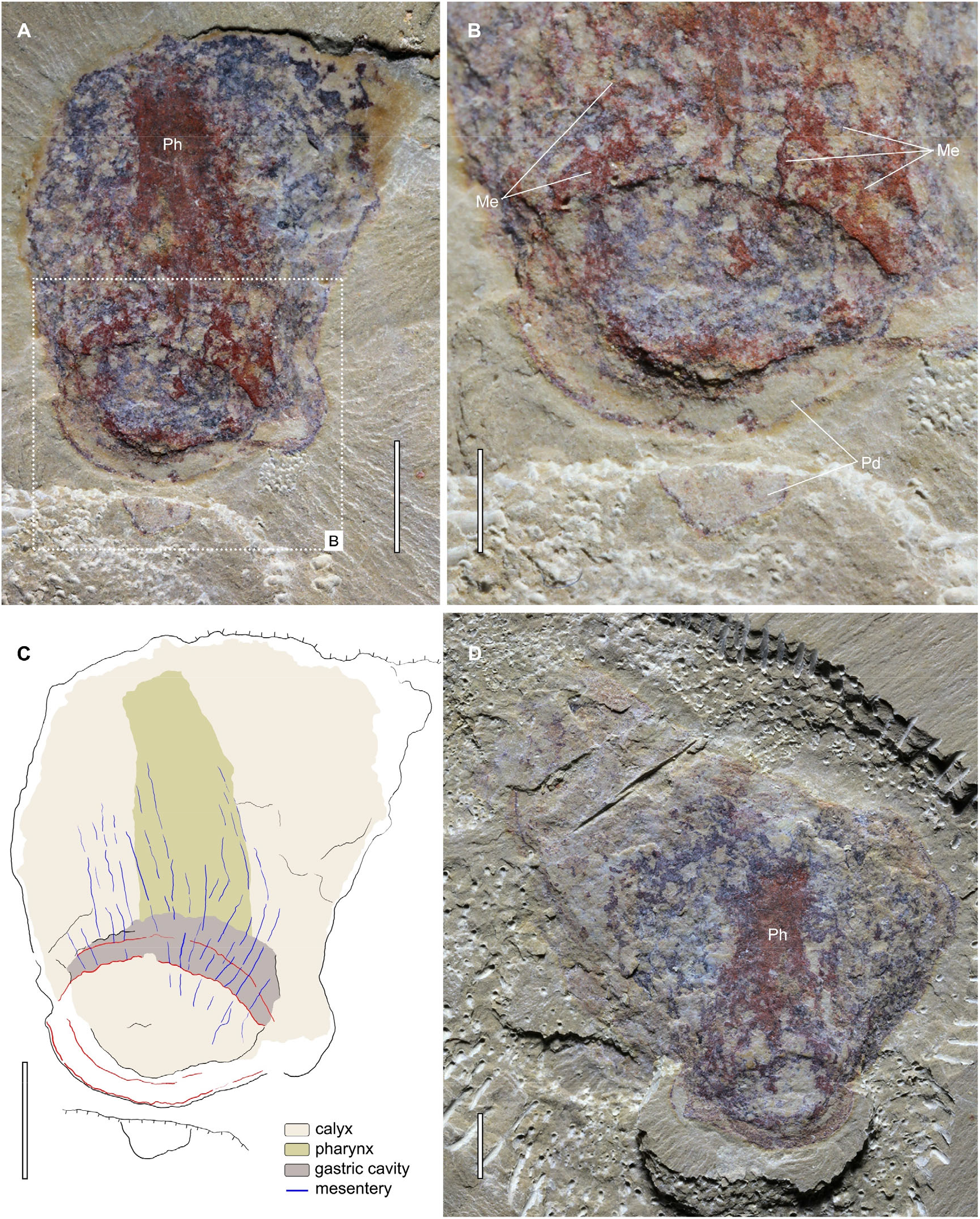
Additional specimen of *C. striata*, preserved in oblique-lateral view. (A-C) YKLP 13492a, (A) an overview of the part, (B) magnification of the body portion, showing a circle-shaped cross section, (C) interpretative drawing of A, red lines draw the outline of the cross section. (D) YKLP 13492b, an overview of the counterpart. Me, mesentery; Pd, periderm; Ph, pharynx. Scale bars: 1 mm (B); 2 mm (A, C, D).

**Fig. S4.**
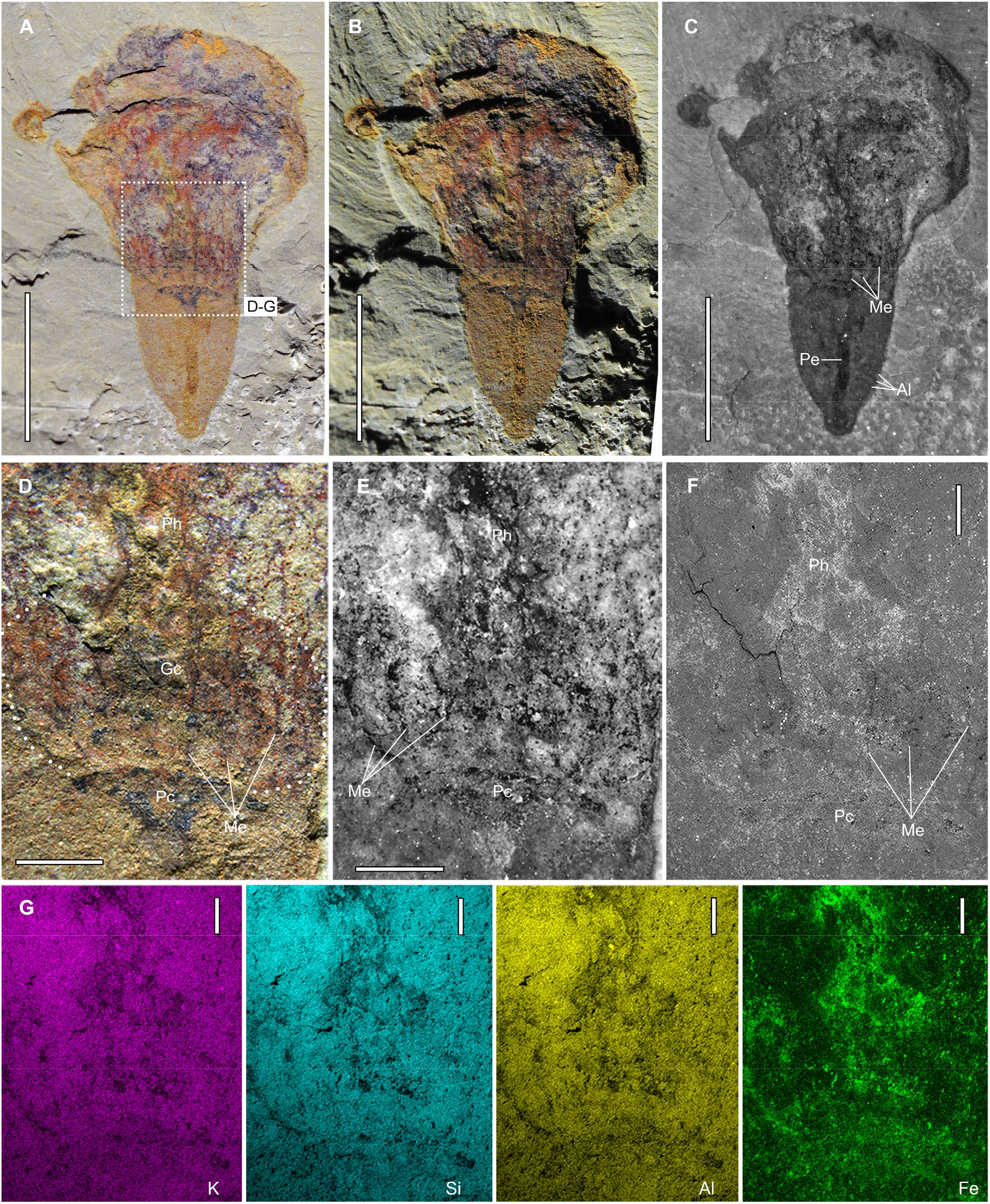
Additional details of *C. striata*, YKLP 13212a (related to Fig. 1A). (A-C) An overview of the part, in high angle light (A), low angle light (B) and fluorescent light (C), respectively. (D-F) Close-up of the cavity region, showing a peduncle chamber, a pharynx and a gastric cavity partitioned by dark lines (mesenteries), in direct light (D), fluorescent light (E) and backscatter of SEM (F), respectively. (G) Elemental maps of the same region as in F, showing high content of iron in the pharynx. Al, annulation; Gc, gastric cavity; Me, mesentery; Pc, peduncle chamber; Pe, peduncle; Ph, pharynx. Scale bars: 500 μm (F and G); 1 mm (D and E); 5 mm (A-C).

**Fig. S5.**
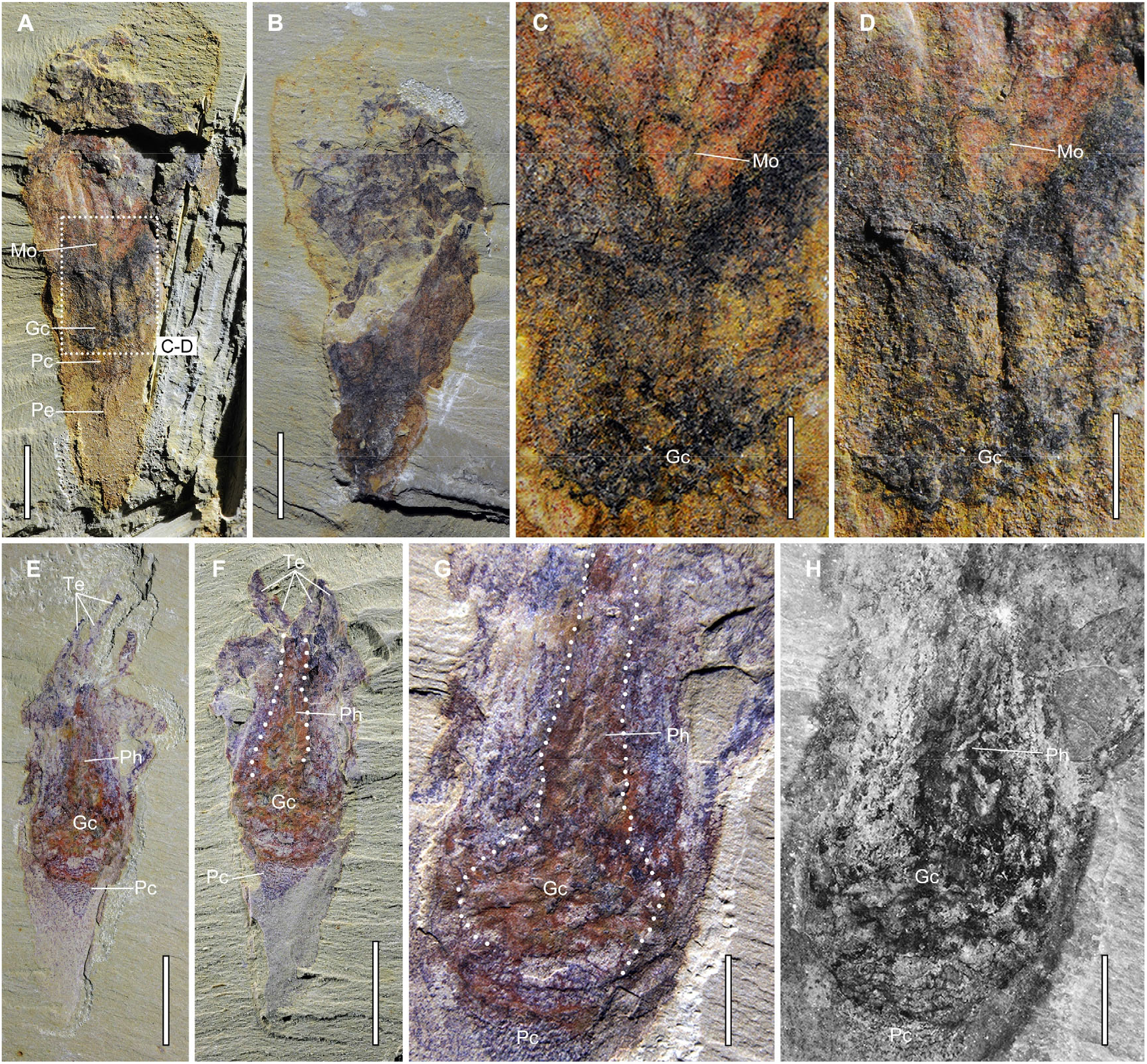
Additional details of *C. striata* (related to Fig. 1D, H). (A) YKLP 13215a, an overview of the part. (B) YKLP 13215b, an overview of the counterpart, showing a globular structure at the distal region. (C-D) Close-up of the pharynx region in A, showing a tongue-shaped dark structure, possible a mouth of the polyp, imaged under high angle light (C) and low angle light (D), respectively. (E) YKLP 13484a, an overview of the part. (F) YKLP 13484b, an overview of the counterpart. (G-H) Close-up of the digestive tract in E, showing a longitudinal pharynx and an expanded gastric cavity, in direct light (G) and fluorescent light (H), respectively. Gc, gastric cavity; Mo, mouth; Pc, peduncle chamber; Pe, peduncle; Ph, pharynx; Te, tentacle. Scale bars: 2 mm (C, D, G, H); 5 mm (A, B, E, F).

**Fig. S6.**
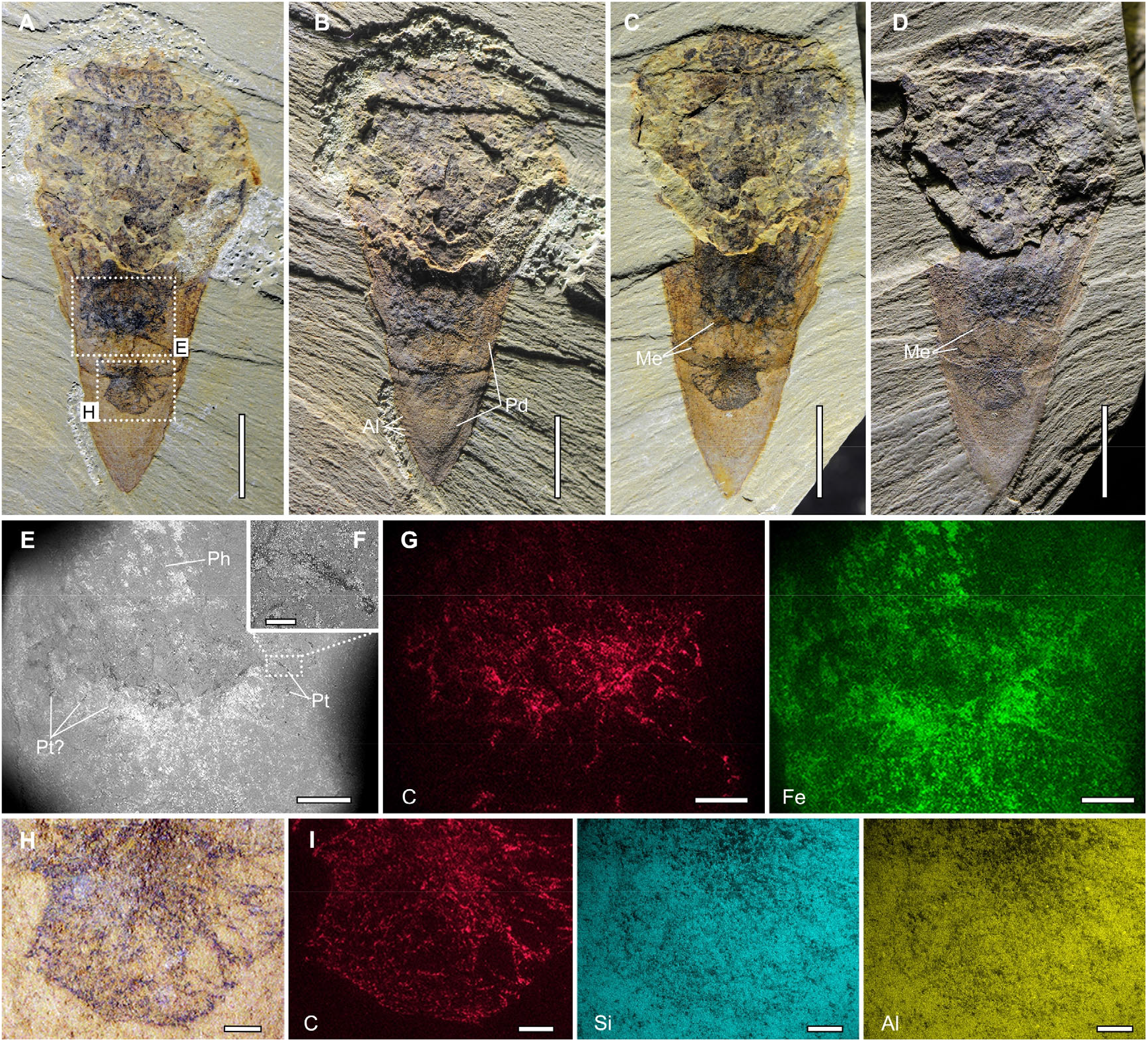
Additional details of *C. striata*, YKLP 13213 (related to Fig. 1C). (A-B) YKLP 13213a, an overview of the part, under high angle light (A) and low angle light (B), respectively. (C-D) YKLP 13213b, an overview of the counterpart, under high angle light (C) and low angle light (D). (E) SEM backscatter of the cavity region, possibly surrounding with peridermal teeth. (F) Close-up of a peridermal tooth. (G) Elemental maps of the same region as in E. (H) Close-up of the peduncle chamber. (I) Elemental maps of the same region of H, showing high content of carbon in the peduncle chamber. Al, annulation; Me, mesentery; Pd, periderm; Ph, pharynx; Pt, peridermal teeth. Scale bars: 100 μm (F); 500 μm (H and I); 1 mm (E and G); 5 mm (A-D).

**Fig. S7.**
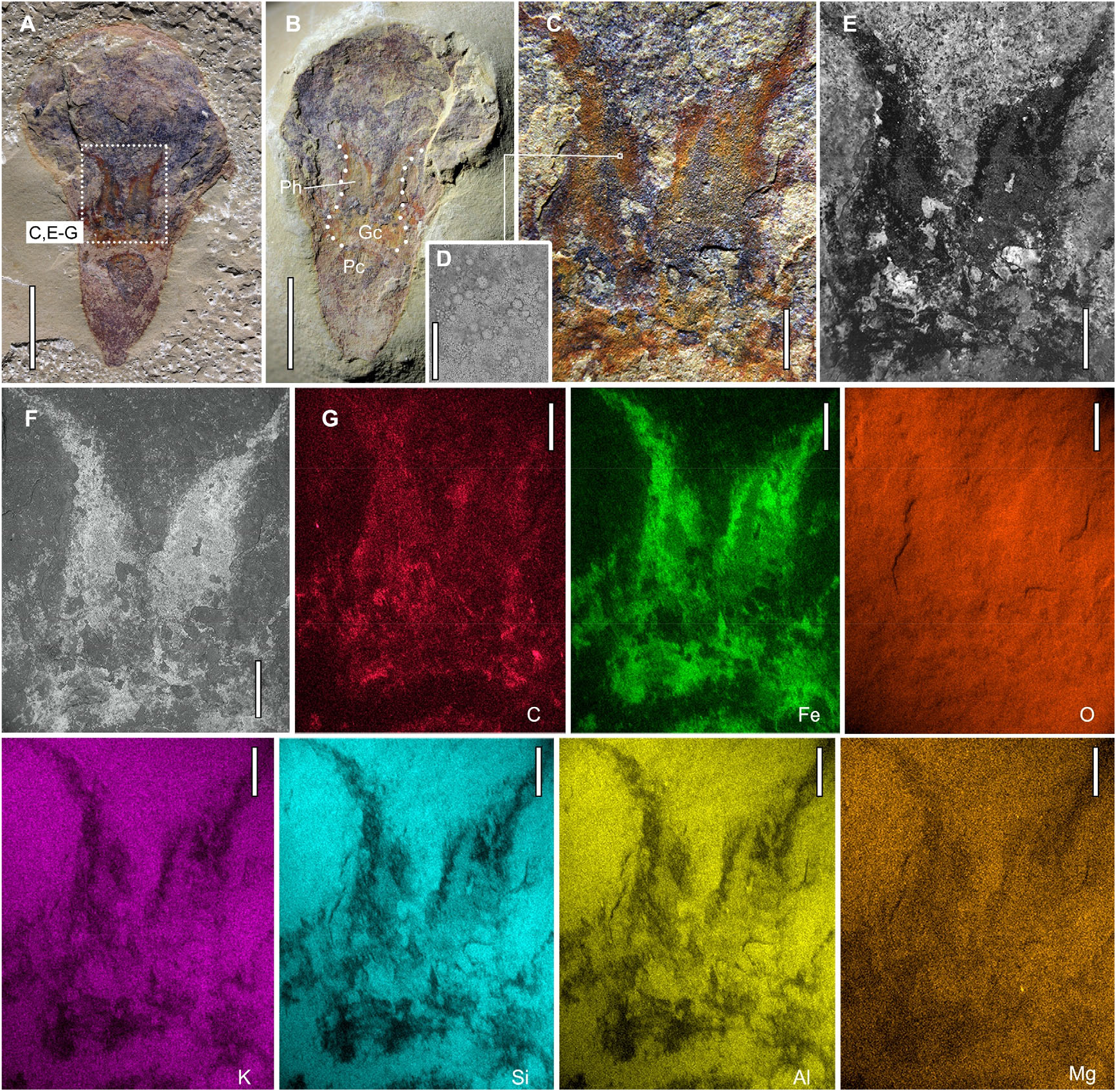
Additional details of *C. striata*, YKLP 13493. (A) YKLP 13493a, an overview of the part. (B) YKLP 13493b, an overview of the counterpart, showing a pharynx, a gastric cavity and a peduncle chamber. (C) Close-up of the pharynx region in A. (D) SEM backscatter of the pharynx in C, showing abundant weathered pyrite granules gathered in the pharynx. (E-F) The same view as in C, under fluorescent light (E) and backscatter of SEM (F), respectively. (G) Elemental maps of the same view as in F, showing high content of carbon and iron in the pharynx. Gc, gastric cavity; Pc, peduncle chamber; Ph, pharynx. Scale bars: 50 μm (D); 1 mm (C, E-G); 5 mm (A and B).

**Fig. S8.**
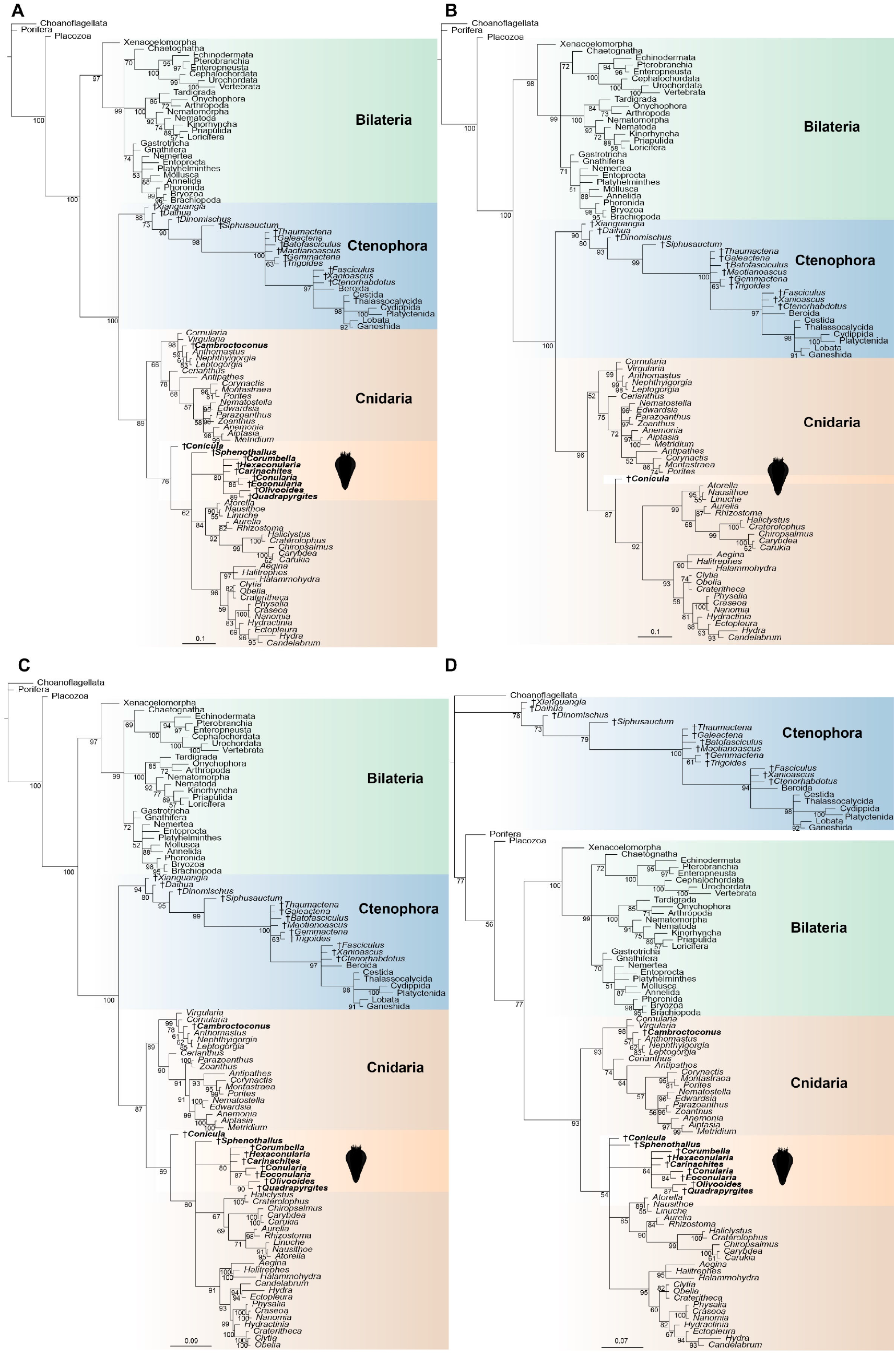
Additional results of phylogenetic analyses (related to Fig. 4C, D). (A) The full results of Fig. 4C, with all the sampled taxa under the condition of unconstraint. (B) The newly added tubular fossils and *Cambroctoconus* are removed, and the topology is unconstrained. *C. striata* is also recovered as a stem group medusozoan. (C) The in-group relationships of cnidarians are constrained based on recent phylogenomic results. *C. striata* is still recovered as a stem group medusozoan. (D) Ctenophores are constrained as the sister group of all other metazoans (ctenophore-first) and cnidarians are constrained as the sister group of bilaterians. *C. striata* is recovered as a total group medusozoan, and is resolved in a clade that is in a polytomy with extant medusozoan taxa and tubular fossils. Numbers at nodes refer to posterior probabilities and scale bars are in units of expected number of substitutions per site. The fossil taxa are indicated by dagger symbol.

**Table S1.**
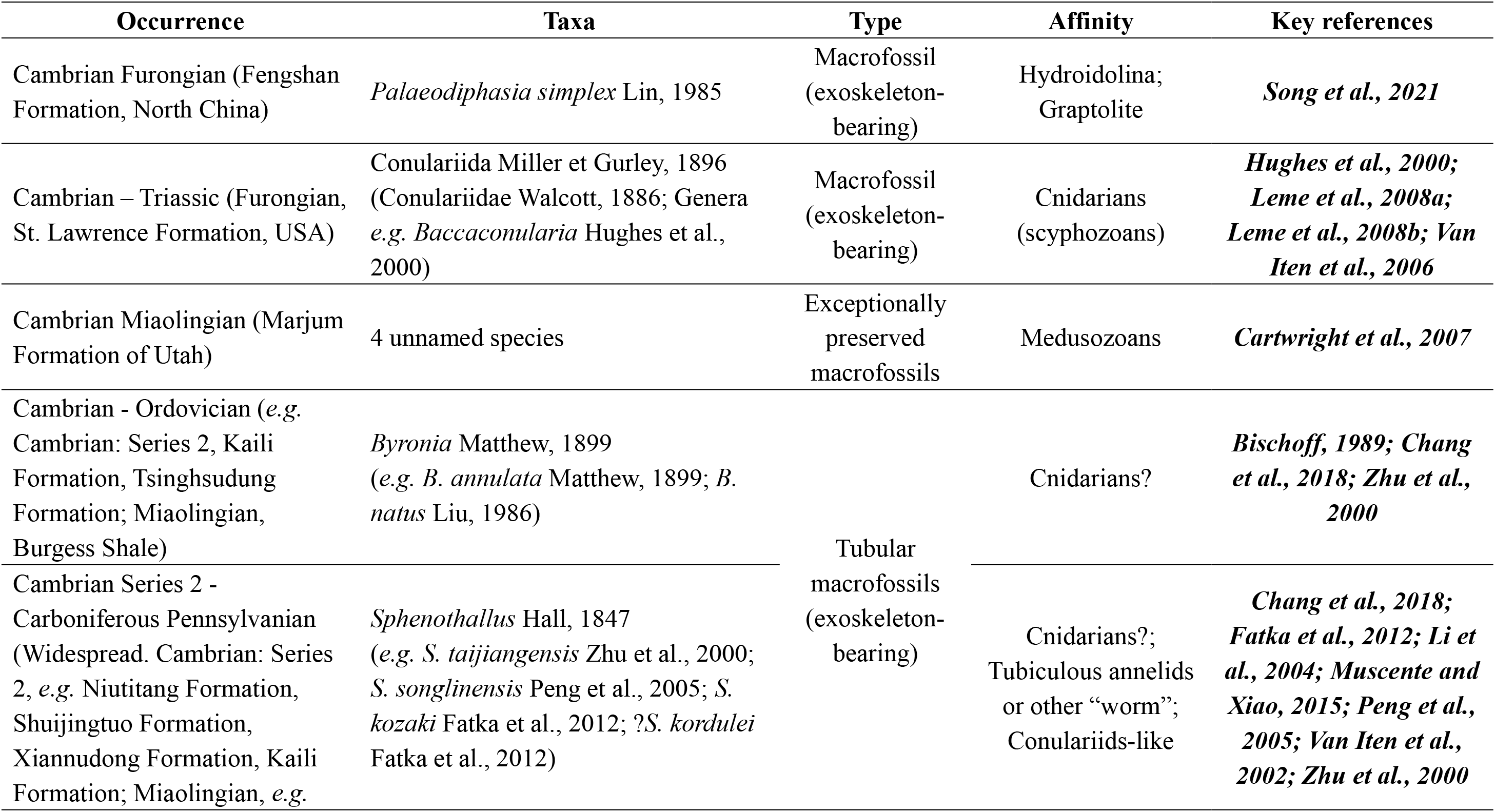
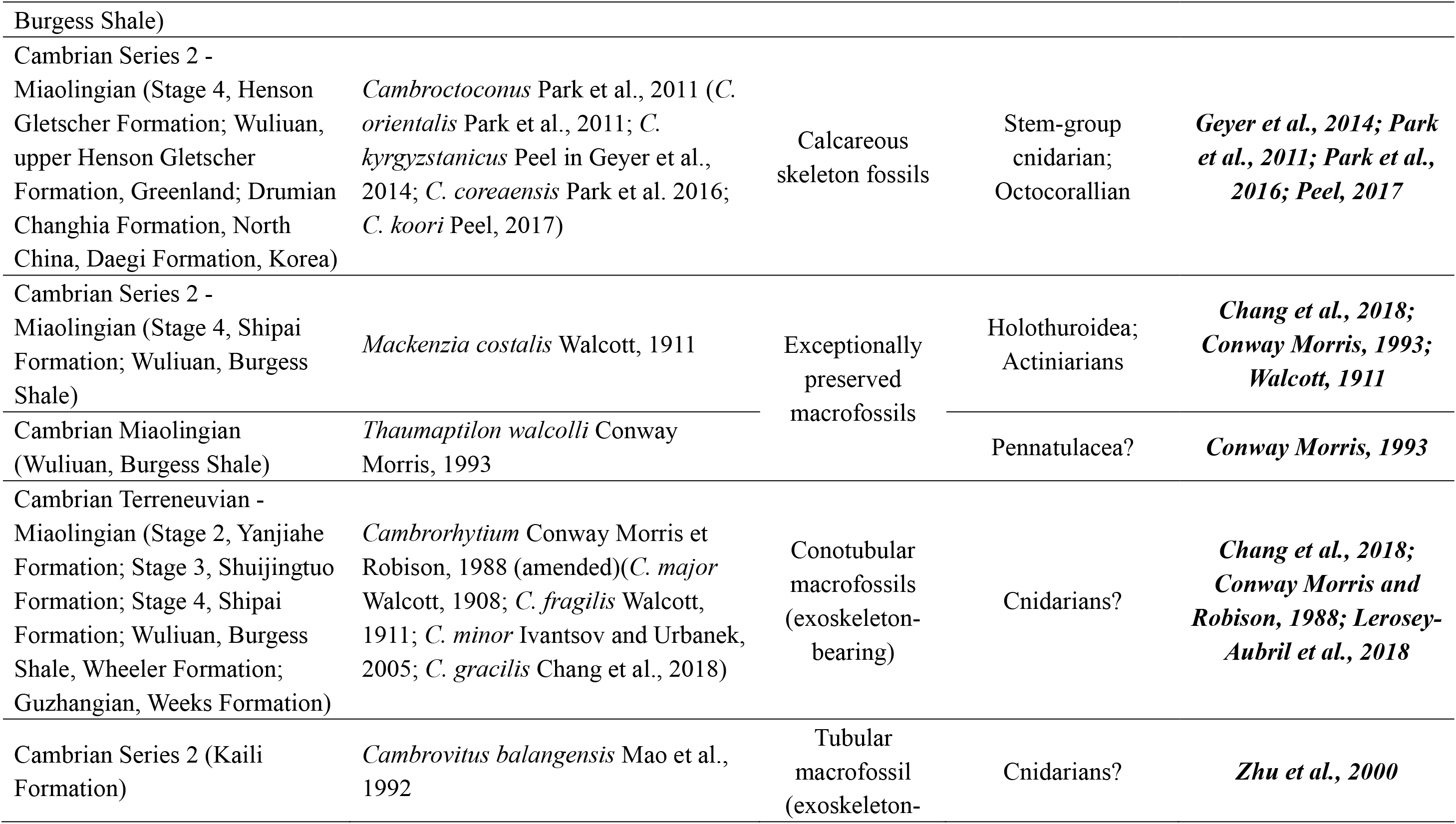
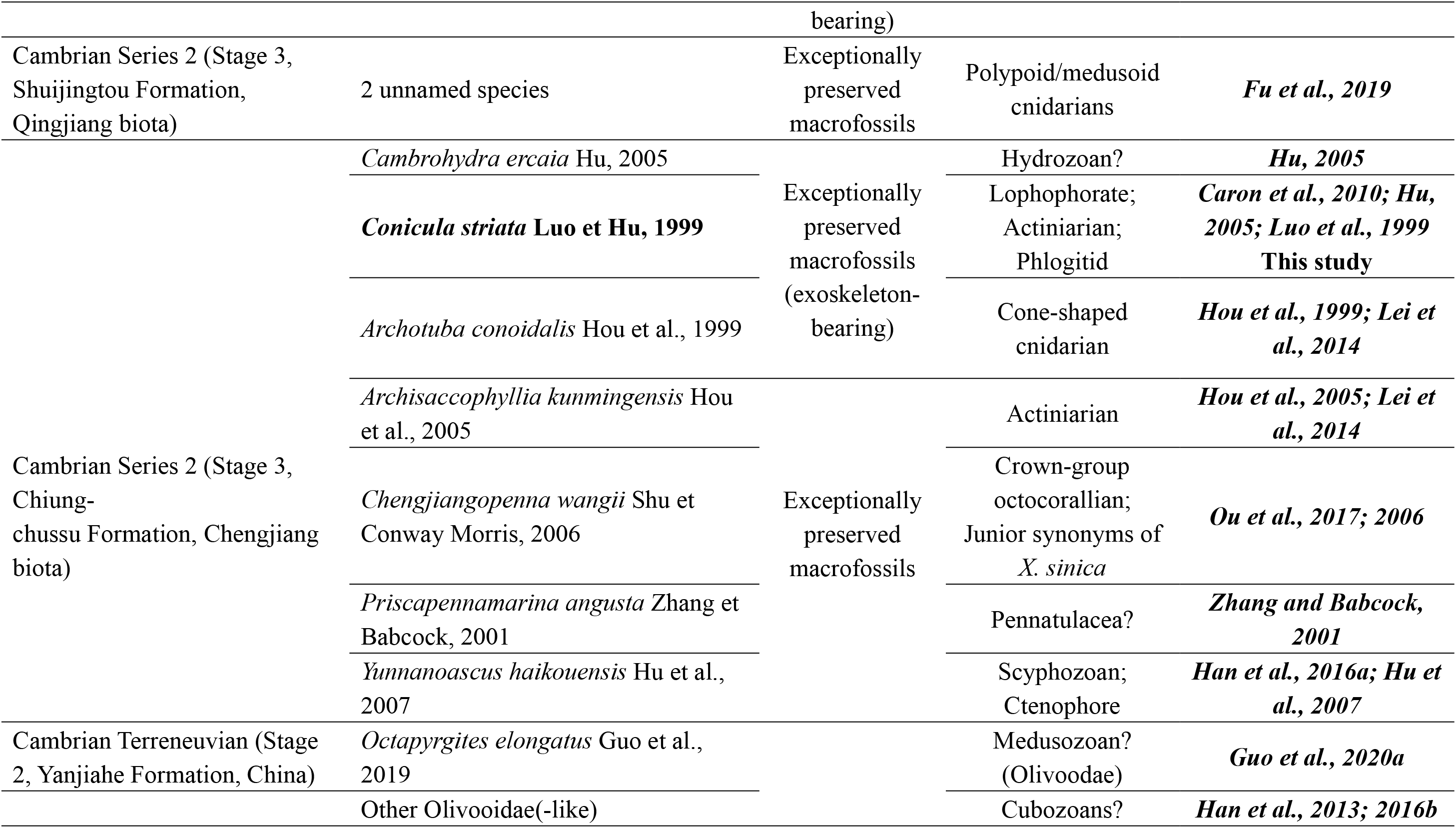
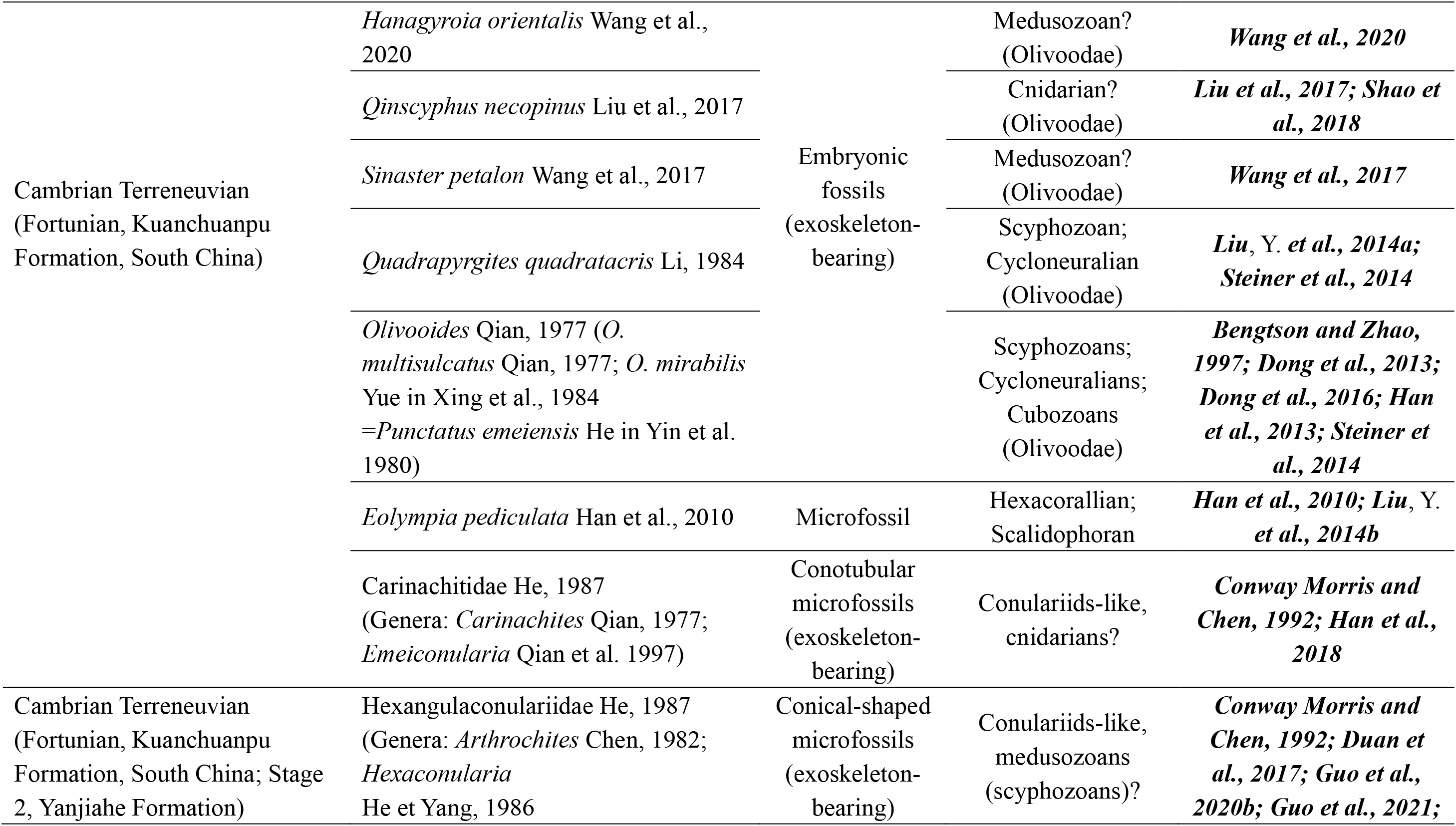
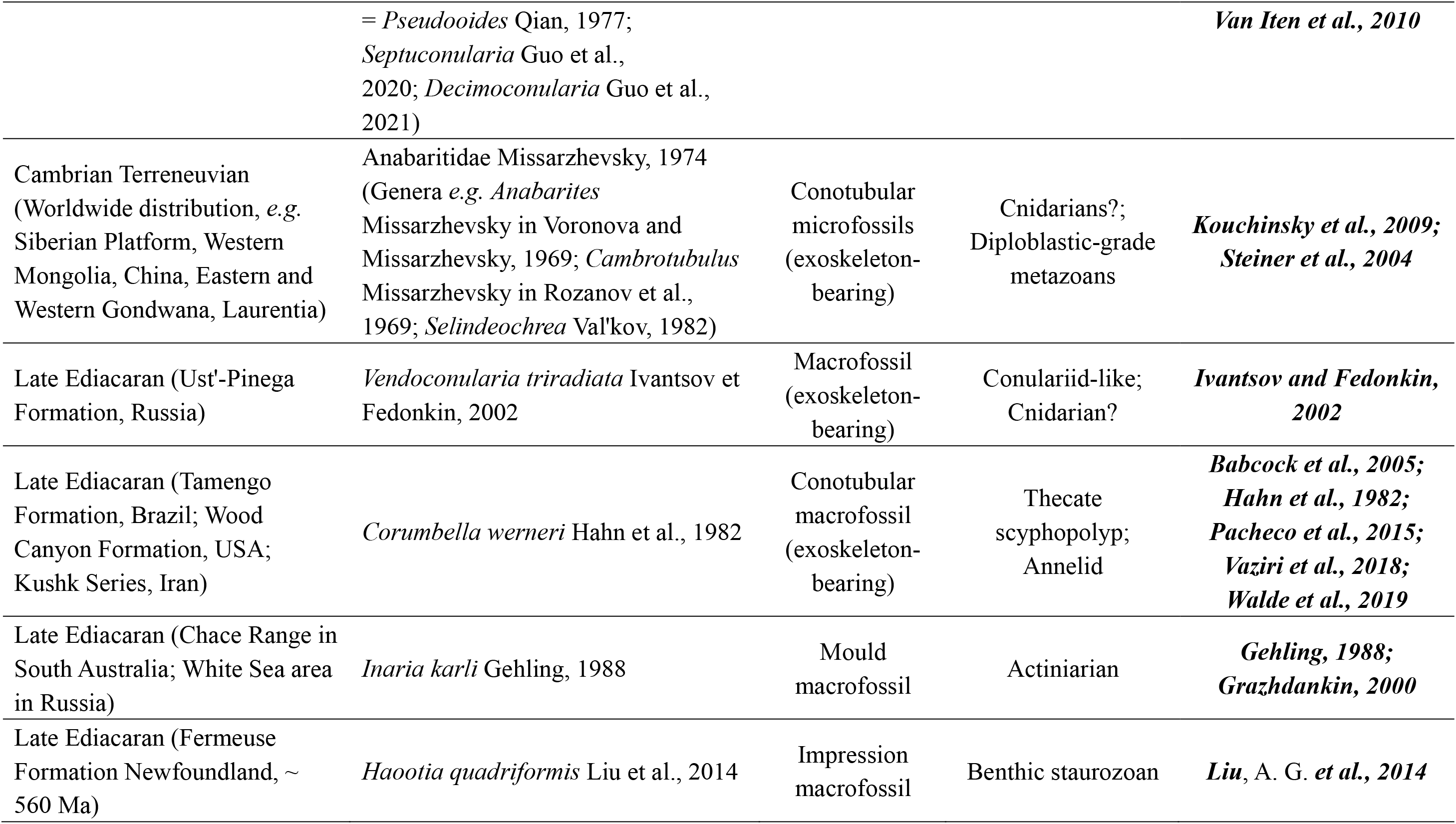
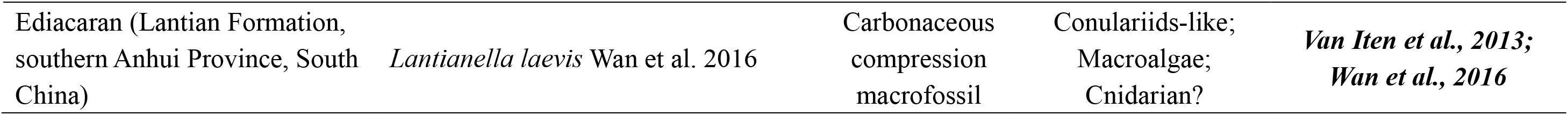
The main occurrence of potential cnidarian fossils in Ediacaran-Cambrian period.

### Phylogenetic information

Our character dataset was mainly adopted from Zhao et al. (Zhao et al., 2019), which includes the main taxa and characters of cnidarians and ctenophores. Our changes to previously existing characters were mainly deleting duplicate or meaningless characters, correcting some mistakes and reordering most of characters. We deleted some redundant taxa (mainly octocorals and siphonophores) as well as *Eolympia* and *Namacalathus* that are useless in current studies. Except for *Conicula*, we also added 16 newly sampled taxa, including extant cnidarian *Edwardsia*, *Zoanthus*, *Craterolophus*, *Carybdea*, *Chiropsalmus*, *Carukia*, *Atorella*, *Linuche*, *Crateritheca* and *Halammohydra*, and tubular fossils *Corumbella*, *Conularia*, *Eoconularia*, *Hexaconularia* and *Carinachites*, and cup-shaped cnidarian fossil *Cambroctoconus*.

The phylogeny analyses were conducted using Bayesian inference under the mkv + gamma model in MrBayes 3.2.7 (Ronquist et al., 2012). We set the number of generations to be 20,000,000 and allowed the stop rule when the average deviation of split frequencies dropped below 0.01, with convergence checked for all parameters (ESS scores >200) using the command ‘sump’. We performed two sets of analyses without any topological constraints (convergence was achieved after <6,000,000 generations in both analyses), one included all sampled taxa and the other removed the newly sampled tubular fossils and *Cambroctoconus*, of which all lack the preservation of reliable soft tissues and therefore have large amounts of missing data (over 80%). However, some recovered clades within the cnidarian have in-group relationships that are inconsistent with the results of recent phylogenomic analyses (Kayal et al., 2018; e.g. McFadden et al., 2021; Zapata et al., 2015). To get further results of the position of *Conicula* in phylogenetic trees, we conducted additional analyses using the command of ‘constraint partial’. All fossil taxa were left unconstrained, so they can wander to any clades in the tree. The first topological constraint was combined with the recent common results of phylogenomic studies in cnidarian clades (Collins et al., 2006; e.g. McFadden et al., 2021). While the second focused on the ctenophore-first (ctenophores are the sister group to all other metazoans including sponges) (e.g. *Dunn et al., 2008; Whelan et al., 2017*) and Planulozoa (for this paper, we only considered cnidarians as the sister group to bilaterians, with the exclusion of Placozoa) (e.g. *Pisani et al., 2015; Simion et al., 2017*). Our commands using for implementing topological constraints in MrBayes 3.2.7 are listed below.

#### Character descriptions

The new characters are in bold type and marked with an asterisk.

1. Collar complex 0 – absent 1 – present
2. Multicellularity with extracellular matrix 0 – absent 1 – present
3. Extracellular digestion 0 – absent 1 – present
4. Ostia with porocytes 0 – absent 1 – present
5. Septate junctions (SJs) 0 – absent 1 – present
6. Tight junctions (TJs) 0 – absent 1 – present
7. Gap junctions (GJs) 0 – absent 1 – present
8. Adherens junctions (AJs) 0 – absent 1 – present
9. Hemidesmosomes 0 – absent 1 – present
10. Epithelia 0 – absent 1 – present
11. Basal laminae 0 – absent 1 – present
12. Collagen 0 – absent 1 – present
13. Nerve cells 0 – absent 1 – present
14. Acetylcholine used as a neurotransmitter 0 – absent 1 – present
15. Diffuse nervous system 0 – absent 1 – present
16. Hox/ParaHox gene 0 – absent 1 – present
17. Epidermis with pulsatite bodies 0 – absent 1 – present
18. Xenacoelomorph cilia 0 – absent 1 – present
19. Striated ciliary rootlets 0 – absent 1 – present
20. Diploblasts made of 2 cell layers 0 – absent 1 – present
21. Triploblasts made of 3 cell layers 0 – absent 1 – present
22. Spiral cleavage with 4d mesoderm 0 – absent 1 – present
23. Through-gut *Conicula* possesses a partitioned, blind gut with only one opening, the type of gut is incompatible with the through-gut bearing two separated openings. Therefore, the character of through-gut is scored as absent in *Conicula*. 0 – absent 1 – present
24. U-shaped gut 0 – absent 1 – present
25. Protonephridia (or homologues) 0 – absent 1 – present
26. Fate of blastopore 0 – protostomy 1 – deuterostomy 2 – amphistomy
27. Body cuticle with chitin 0 – absent 1 – present
28. Body cuticle with α-chitin 0 – absent 1 – present
29. Body cuticle moulted 0 – absent 1 – present
30. Segmented body with jointed limbs 0 – absent 1 – present
31. Lobopods 0 – absent 1 – present
32. Slime papillae 0 – absent 1 – present
33. Telescoping mouth cone with protrudable stylets 0 – absent 1 – present
34. Respiration via metameric tracheae and spiracles 0 – absent 1 – present
35. Teloblastic segmentation 0 – absent 1 – present
36. Longitudinal ventral nerve cord(s) 0 – absent 1 – present
37. Circum-pharyngeal, collar-shaped brain with anterior and posterior rings of perikarya separated by a ring-shaped neuropil 0 – absent 1 – present
38. Introvert with scalid rings 0 – absent 1 – present
39. Flosculi 0 – absent 1 – present
40. Immunoreactivity of horseradish peroxidase (HRP) 0 – absent 1 – present
41. Lophophore 0 – absent 1 – present
42. Trochophore 0 – absent 1 – present
43. Radula 0 – absent 1 – present
44. Segmental metanephridia sacculus 0 – absent 1 – present
45. Chitinous microvillar appendages (chaetae) 0 – absent 1 – present
46. Parapodia with dorsal and ventral branches terminated by β-chitinous chaetae 0 – absent 1 – present
47. Eversible proboscis surrounded by rhynchocoel 0 – absent 1 – present
48. Complex jaw apparatus in pharynx 0 – absent 1 – present
49. Mesoderm 0 – absent 1 – present
50. Origin of mesoderm 0 – from the blastopore lips and as ectomesoderm 1 – from the walls of the archenteron or neural crest
51. Mixocoel (haemocoel) surrounded by segmented mesoderm 0 – absent 1 – present
52. Radial cleavage 0 – absent 1 – present
53. Coelom formation 0 –schizocoely 1 –enterocoely
54. Trimeric coelom 0 – absent 1 – present
55. Pharyngeal slits 0 – absent 1 – present
56. Endostyle (or homologues) 0 – absent 1 – present
57. Notochord 0 – absent 1 – present
58. Stomochord 0 – absent 1 – present
59. Haemal system with axial complex 0 – absent 1 – present
60. Calcareous endoskeleton composed of separate ossicles 0 – absent 1 – present
61. Tornaria type larva 0 – absent 1 – present
62. Longitudinal dorsal nerve cord 0 – absent 1 – present
63. Zig zag myomeres 0 – absent 1 – present
64. Endothelium that lines the inner wall of blood vessels 0 – absent 1 – present
65. Neural crest 0 – absent 1 – present
66. Neurogenic placodes 0 – absent 1 – present
67. Dorsoventral axis 0 – absent 1 – present
68. Anterior posterior axis 0 – absent 1 – present
69. **Symmetry\*** The single character of symmetry in Zhao et al. (character 69)(Zhao et al., 2019) is now split into two characters (69 and 70) to establish character polarity, as scoring the different types of symmetry would necessitate scoring the character of symmetry as absent for Choanoflagellata and Placozoa. The character is here scored as polymorphic for Porifera because the symmetry is present only in some subgroups of sponges, such as calcareous sponges (Manuel, 2009) or Palaeozoic sponges (Botting et al., 2014). The material of *Conicula* exhibits a circular cross-section, along with the conical gross morphology (with oral-aboral axis), suggesting the presence of symmetric nature. 0 – absent 1 – present
70. Type of symmetry We are careful to score this character as uncertain in *Conicula* because the total number of tentacles or mesenteries in current material is yet to be determined. 0 – bilateral 1 – biradial 2 – triradial 3 – tetraradial 4 – pentaradial 5 – hexaradial
71. Compression in pharyngeal plane 0 – absent 1 – present
72. Compression in oral aboral axis 0 – absent 1 – present
73. Compression in tentacular plane 0 – absent 1 – present
74. Cydippid larvae 0 – absent 1 – present
75. Ciliary rosettes 0 – absent 1 – present
76. Radially-arranged outgrowths from the interface between the oral and aboral regions *Conicula* possesses soft outgrowths (tentacles) extending from the oral disc of the polyp, so this character is scored as present in *Conicula*. 0 – absent 1 – present
77. Radial outgrowths fixed in globular configuration 0 – absent 1 – present
78. Radial outgrowths tentacular 0 – absent 1 – present
79. Outgrowths with pinnules The tentacles in *Conicula* are unbranched with no signs of the existence of pinnules. We also code this character as inapplicable in those taxa without tentacular outgrowth because the possession of pinnules is contingent on the presence of tentacles. 0 – absent 1 – present
80. Outer sheaths on external surface of radial outgrowths This character is coded as absent in *Conicula* because no evidence of outer sheaths is present in the external surface of the outgrowths in *Conicula*. 0 – absent 1 – present
81. Outgrowths with ciliary rows 0 – absent 1 – present
82. Ciliary rows paired 0 – paired 1 – unpaired
83. Orientation of ciliary rows relative to oral-aboral axis 0 – adaxial 1 – abaxial
84. Uniformity of ciliary rows 0 – uniform 1 – non uniform
85. Number of ciliary rows 0 – eight 1 – eighteen 2 – six 3 – twenty four 4 – more than 24 5 – sixteen
86. Cushion rings/plates or polster cells 0 – absent 1 – present
87. Cushion rings paired 0 – paired 1 – unpaired
88. Large compound cilia 0 – absent 1 – present
89. Large cilia fused to form locomotory plate 0 – absent 1 – present
90. Extension of the oral surface to form oral cone 0 – absent 1 – present
91. Aboral region represented only by apical organ 0 – absent 1 – present
92. Apical organ forming narrow pointed extension 0 – absent 1 – present
93. Oral macrocilia 0 – absent 1 – present
94. Oral lobes 0 – absent 1 – present
95. Morphology of tip of oral extension 0 – narrow 1 – voluminous 2 – manubrium-like
96. Mouth as margin of creeping sole 0 – absent 1 – present
97. Pharyngeal ridges 0 – absent 1 – present
98. Sealing ridges of pharynx 0 – absent 1 – present
99. Macrocilia on pharynx lining inside of mouth 0 – absent 1 – present
100. Ciliary dome 0 – absent 1 – present
101. Statolith 0 – absent 1 – present
102. Balancers 0 – absent 1 – present
103. Pole plates 0 – absent 1 – present
104. Aboral papillae 0 – absent 1 – present
105. Ciliary grooves 0 – absent 1 – present
106. Interplate ciliary groove (ICG) 0 – absent 1 – present
107. Pharyngeal canals 0 – absent 1 – present
108. Tentacular canals 0 – absent 1 – present
109. Meridional canals 0 – absent 1 – present
110. Diverticula of meridional canals 0 – absent 1 – present
111. Circumoral ring canal 0 – absent 1 – present
112. Termination of meridional and pharyngeal canals 0 – both terminate blindly 1 – branch to form a complex network 2 – united with the circumoral ring canal
113. Interradial canals 0 – absent 1 – present
114. **Number of interradial canals\*** This character ‘interradial canals’ in the previous matrix combined the presence and number of interradial canals (Zhao et al., 2019), and is now split into two separate characters. 0 – two 1 – four
115. Adradial canals 0 – directly branch from the infundibulum 1 – branch from interradial canals
116. Aboral canal 0 – absent 1 – present
117. Anal canals 0 – absent 1 – present
118. Anal pores 0 – absent 1 – present
119. Paired ctenophore Tentacles 0 – absent 1 – present
120. Colloblasts 0 – absent 1 – present
121. Tentilla 0 – absent 1 – present
122. Disposition of tentilla 0 – fringing along tentacles 1 – fringing along elongate oral margin
123. Tentacle sheaths 0 – absent 1 – present
124. Opening position of tentacles 0 – orally 1 – aborally
125. Auricles 0 – absent 1 – present
126. Brood chambers 0 – absent 1 – present
127. Sclerotised arms and calyx 0 – absent 1 – present
128. Tentacles extending beyond skeletal rods and outer sheaths 0 – absent 1 – present
129. Medial structures of skeletal elements 0 – absent 1 – present
130. Kinked spokes 0 – absent 1 – present
131. Spinose spokes 0 – absent 1 – present
132. Radiating flaps/lobes 0 – absent 1 – present
133. Oral structure separated by circumferential constriction 0 – absent 1 – present
134. Constriction type 0 – skirt or bell 1 – lappets
135. Feeding strategy in tentaculate metazoans Metazoans who rely on pinnules/cilia to feeding are defined as the suspension-feeding groups, of which feeding strategies are mainly the granules suspended in the water current. Those metazoans bearing flexible, non-cilia tentacles are usually preying on macro-organisms. *Conicula* has flexible, free bending tentacles, with no clear signs of pinnules/cilia, it is coded as predominantly macro for feeding strategy. 0 – predominantly micro 1 – predominantly macro
136. Mesoglea 0 – absent 1 – present
137. Mesoglea cellular 0 – absent 1 – present
138. Embryonic development 0 – direct 1 – indirect
139. Structure of mitochondrial DNA 0 – circular 1 – linear
140. Cnidae 0 – absent 1 – present
141. Cnidae in gastrodermis 0 – absent 1 – present
142. Cnidocil 0 – absent 1 – present
143. **Cnidae apical structure\*** The typical feature of anthozoan cnidae is the absence of operculum (Brusca et al., 2016), and we score the sampled taxa mainly following Reft & Daly (Reft and Daly, 2012). 0 – flaps/cap 1 – operculum 2 – no flaps/operculum
144. Spirocyst 0 – absent 1 – present
145. Ptychocyst 0 – absent 1 – present
146. Stenoteles 0 – absent 1 – present
147. Euryteles 0 – absent 1 – present
148. Birhopaloids 0 – absent 1 – present
149. Rhopalonemes 0 – absent 1 – present
150. Desmonemes 0 – absent 1 – present
151. Mastigophores 0 – absent 1 – present
152. Isorhizas 0 – absent 1 – present
153. Basitrichous isorhizas 0 – absent 1 – present
154. Apotrichous isorhizas 0 – absent 1 – present
155. Heterotrichous anisorhizas 0 – absent 1 – present
156. **Nemathybomes on scapus\*** *Edwardsia* is characterised by prominent nemathybomes that is nematocyst-dense pockets in the epidermis, which is absent in *Nematostella* (Daly, 2002). It is inapplicable in other cnidarians as their body column is not composed of scapus. 0 – absent 1 – present
157. Zooxanthellae 0 – absent 1 – present
158. Mesogleal skeleton 0 – absent 1 – present
159. Ectodermal skeleton 0 – absent 1 – present
160. Composition of ectodermal skeleton 0 – proteinaceous 1 – calcitic
161. **Corallum\*** Corallum is a spiny, proteinaceous skeleton type widely present in Antipatharia (Daly et al., 2007). 0 – absent 1 – present
162. Columella 0 – absent 1 – present
163. Costae 0 – absent 1 – present
164. Octocorallian spicules 0 – absent 1 – present
165. Spicules in tentacle 0 – absent 1 – present
166. Gorgonin 0 – absent 1 – present
167. Periderm Considering the chemical composition varies in different types of periderm based on Mendoza-Becerril et al. (Mendoza-Becerril et al., 2016), we also code the periderm as present in *Hydra*. *Cornularia* is a unique genus among Octocorallians in having a polyp covered by a theca-like chitinous outer sheath (López-González et al., 1995; Weinberg, 1978), we score *Cornularia* as present for a periderm as well. The conical external skeleton of *Conicula* is a robust structure ornamented by parallel annulations, encasing fully the internal polyp. These features are consistent with the tubular fossils with an external periderm, such as conulariids. We therefore code periderm as present in *Conicula*. 0 – absent 1 – present
168. Periderm type Based on Mendoza-Becerril et al., the periderm is assigned into two types of corneous (chitin-protein) and coriaceous (calcium carbonate/phosphate), the former is widely present in extant organisms, while the latter is thought to be present in a few of Anthoathecata, such as *Millepora*, and most of fossil groups, like conulariids and *Corumbella* (Mendoza-Becerril et al., 2016). *Cornularia* is coded as having the corneous type of periderm because of the possession of a chitinous envelope (López-González et al., 1995; Weinberg, 1978). It is unknown the peridermal type of *Conicula* based on current evidence. 0 – corneous 1 – coriaceous 2 – fibrous
169. Cuticle layers in periderm Conulariids had found evidence of two-layer cuticle present in the periderm (Jerre, 1994; Van Iten, 1992a). *Corumbella* may have one-layer cuticle in the periderm (Mendoza-Becerril et al., 2016). While the number of cuticle layers in *Conicula* is uncertain. 0 – one 1 – two 2 – five
170. **Regions with periderm\*** A new character describes the body region encased by the periderm and is scored mainly following Mendoza-Becerril et al. (Mendoza-Becerril et al., 2016). Basing on fossil material, *Conicula* and other tubular fossils have a periderm encasing the entire polyp. 0 – entire polyp 1 – hydrocaulus or hydrorhiza 2 – basal region or podocyst 3 – stolon and anthostele
171. **Tubular periderm\*** A new character defines the shape of periderm. The tubular periderm is present in most of extant taxa that have a periderm encasing the entire polyp or hydrocaulus. It also appears in *Conicula* and other tubular fossils, including conulariids, olivooids, *Hexaconularia*, *Carinachites*, *Corumbella* and *Sphenothallus*. 0 – absent 1 – present
172. **Shape of tubular periderm\*** Coronates, *Conicula* and tubular fossils have a cone-shaped periderm, while hydrozoans have two types of tubular periderm shape that are contingent on the encased region of the periderm. 0 – cone 1 – tube 2 – goblet-like
173. **Cone slender and elongate\*** Coronates possess a slender and elongate cone that differs from the pyramid-shaped or conical fossil tubes (such as conulariids and *Conicula*), except for the tubes of *Sphenothallus* (Dzik et al., 2017) and *Corumbella* (Pacheco et al., 2015) that appear to slender and elongate. 0 – absent 1 – present
174. **Tapering end of cone\*** There is a disc-like attachment structure present in the tapering end of the cone of coronates, similar structure could be also seen in the cone of *Sphenothallus* (Dzik et al., 2017). *Conicula*, olivooids (Dong et al., 2016; Liu, Y. et al., 2014a), *Hexaconularia* (Van Iten et al., 2010) and *Carinachites* (Han et al., 2018) lack the structure of attachment disc but instead exhibit a blunt tapering end. It is uncertain that conulariids have an attachment disc, because it is usually broken off at the tapering end of fossil specimens. 0 – blunt 1 – with an attachment disc
175. **Periderm forming a globular expansion\*** This new character defines the periderm shape of *Conicula*. All fossil and living taxa with a tubular periderm do not expand the tube to form a globular chamber in the distal region, and this character is scored as absent in them. 0 – absent 1 – present
176. **Periderm with face\*** Unlike the periderm of extant taxa, the periderm is divided into various numbers of faces in conulariids, olivooids, *Hexaconularia* and *Carinachites*. We follow the most common opinion to code the peridermal face as present in *Corumbella* (Babcock et al., 2005; Pacheco et al., 2011; Pacheco et al., 2015; Van Iten et al., 2016). This character is absent in *Conicula* and *Sphenothallus* since there are no evident faces present in fossil material. 0 – absent 1 – present
177. **Face divided by corner groove\*** This character is generally present in conulariids (Leme et al., 2008b), *Hexaconularia* (Van Iten et al., 2010) and *Carinachites* (Conway Morris and Chen, 1992), but is absent in *Olivooides* (divided by longitudinal ridges) and *Corumbella*. 0 – absent 1 – present
178. **Corner groove wide and deep\*** *Carinachites* possesses wide and deep grooves with triangular shape in cross-section (Han et al., 2018), which are distinct from the narrow and shallow grooves that appear in conulariids and *Hexaconularia*. 0 – absent 1 – present
179. **Midline\*** The character of midline is widely present in the middle of the face of conulariids (Van Iten, 1992b), but is absent in olivooids, *Hexaconularia* and *Carinachites*. We score this character as present in *Corumbella* following the common opinion (Babcock et al., 2005; Pacheco et al., 2011; Pacheco et al., 2015; Van Iten et al., 2016). 0 – absent 1 – present
180. **Carinae\*** This character is widely present in the internal periderm of conulariids (Van Iten, 1992b), and is absent in *Corumbella* following the recent study (Pacheco et al., 2015). 0 – absent 1 – present
181. **Bipartite periderm stage\*** This character is adopted from character 203 ‘embryonic stage retained in the tube morphology’ in previous matrix (Zhao et al., 2019). The tubular periderms of olivooids and *Hexaconularia* are consisting of two distinct body parts, the embryonic tissue and the post-embryonic tissue (Steiner et al., 2014; Van Iten et al., 2010). Extant cnidarians experience a planula phase before forming a polypoid or medusoid body, without a bipartite periderm stage at early development phase. We score this character as absent in extant cnidarians with a tubicolous periderm and as unknown in all other tubular fossils. 0 – absent 1 – present
182. **Apical region with longitudinal ridge\*** Olivooids have longitudinal ridges in the apical cone (embryonic tissue) that play a crucial role to determine the symmetric type (Steiner et al., 2014). *Hexaconularia* appears to with no evident ridges in the apical region (Duan et al., 2017). 0 – absent 1 – present
183. **Number of ridges\*** *Olivooides* has five ridges in the apical cone (Dong et al., 2016), while *Quadrapyrgites* exhibits four ridges (Liu, Y. et al., 2014a). 0 – four 1 – five
184. **Periderm apertural end\*** This new character describes differences of the apertural end of periderm across sampled taxa. The distal portion of periderm tube protrudes towards the central lumen to form apertural lobes surrounding the terminal opening, as seen in conulariids (Moore, 1956), olivooids (Dong et al., 2016; Liu, Y. et al., 2014a) and *Carinachites* (Han et al., 2018). The operculum appeared in the periderm tube of hydrozoan (Bouillon et al., 2006) and coronates (Werner, 1973). *Sphenothallus* (Chang et al., 2018; Dzik et al., 2017) and *Corumbella* (Babcock et al., 2005; Pacheco et al., 2015) do not develop an apertural operculum or oral lobes. *Conicula* and *Hexaconularia* are scored as unknown as it is uncertain based on current fossil material. 0 – open 1 – with lobes 2 – with an operculum
185. **Cross-section of the terminal end of periderm\*** This character is adopted from character 1 of Leme et al. (Leme et al., 2008a). The circle-shaped cross-section of the terminal end of periderm is present in *Conicula*, *Sphenothallus* and extant hydrozoans and coronates. In contrast, conulariids, olivooids, *Hexaconularia*, *Carinachites* and *Corumbella* have various types of polygonal cross-sections. 0 – circular 1 – polygon
186. Peridermal tooth This character is modified based on the code in Zhao et al. (Zhao et al., 2019). Peridermal teeth refer to the inward protrusion of the inner layer periderm to form ridge-like or tooth-like structures towards the lumen. We now score the peridermal tooth as inapplicable in the taxa without a periderm. Several hydrozoans possess intrathecal septa or ridges within the periderm that is presumably homologous to the teeth or ridges of the periderm tube. We hereby add a new taxa *Crateritheca* with prominent intrathecal septa (Bouillon et al., 2006; Millard, 1975) into the sampled taxa. 0 – absent 1 – present
187. **Tooth morphology\*** *Olivooides* has peridermal teeth in the form of paired projections (Dong et al., 2016). *Sphenothallus* possesses simple sheet-like cusps (ridges) with smooth rim (Dzik et al., 2017), which may present in *Eoconularia* (Jerre, 1994) and *Conicula* as well. We also code *Crateritheca* as having sheet-like cusps in the perisarc (Bouillon et al., 2006; Millard, 1975).Coronates bear complex peridermal teeth with various shapes (Jarms, 1991; Jarms et al., 2002). 0 – paired projections 1 – sheet-like cusps (ridges) 2 – teeth-like cusps
188. **Tooth disposition\*** *Sphenothallus* (Dzik et al., 2017), *Eoconularia* (Jerre, 1994) and extant *Crateritheca* (Millard, 1975) have irregular arrangements of internal ridges. *Conicula*, *Olivooides* (Dong et al., 2016) and extant coronates (Jarms, 1991; Jarms et al., 2002) have peridermal teeth arranged in the whorls. 0 – irregular arrangement 1 – whorl
189. **Periderm with annulation\*** The external annulation in the periderm is widely present in the tubular fossil groups, including conulariids, olivooids, *Corumbella*, *Carinachites* and *Conicula*, as well as in extant coronates and most hydrozoans. Most tubes of *Sphenothallus* do not have obvious annulations, but some of them exhibit faint, fine striae (Chang et al., 2018; Muscente and Xiao, 2015; Zhu et al., 2000), we also code annulation as present in the tube of *Sphenothallus*. 0 – absent 1 – present
190. **Annulation distribution\*** Most external annulations in the periderms of hydrozoans are confined to the basal portion, and this phenomenon appears also in *Conicula*. We score the external annulations in coronates, conulariids, olivooids, *Sphenothallus*, *Corumbella* and *Carinachites* as widespread in the periderm tube. 0 – confined to basal part 1 – widespread in periderm
191. **Annulation continuous** The annulations in extant coronates and hydrozoans are continuous around the entire periderm. We score the continuous annulation as present also in *Conicula*, *Sphenothallus* and olivooids since they do not have external midlines or corner grooves. 0 – absent 1 – present
192. **Location of annulation offset\*** Conulariids are coded following characters 5 and 6 of Leme et al. (Leme et al., 2008a). The annulations of *Corumbella* (Pacheco et al., 2015) and *Hexaconularia* (Van Iten et al., 2010) offset in the midlines, and the annulations are interrupted in the corner sulcus of *Carinachites* (Conway Morris and Chen, 1992). 0 – interradii (midlines) 1 – perradii (corners)
193. **Annulation type\*** Based on the morphology of external annulations preserved in the fossil material, we score olivooids as the special crest-like annulations (Dong et al., 2016; Liu, Y. et al., 2014a), while living hydrozoans and coronates as well as *Conicula* and other tubicolous periderms (Babcock, 1991) exhibit the rib-like annulations, except for *Sphenothallus*, which has faint, fine striae that distinctly differ from the rib-like annulations (Chang et al., 2018; Zhu et al., 2000). 0 – rib-like 1 – crest-like 2 – striae
194. **Ornament in rib\*** The external annulations in extant coronates and hydrozoans are smooth without ornaments, which are also applicable in *Conicula*, *Sphenothallus*, *Carinachites* and *Corumbella*. The ornaments in ribs of conulariids are variable across different taxa (Leme et al., 2008a). *Eoconularia* does not have ornaments in the transverse ribs, while *Conularia* has nodes in the transverse ribs (Babcock, 1991; Leme et al., 2008a). 0 – absent 1 – present
195. Propagation through lateral budding 0 – absent 1 – present
196. Oocyte development 0 – oocytes develop without accessory cells 1 – oocytes develop with accessory cells 2 – oocytes develop within follicles 3 – oocytes develop from uptake of somatic or other germ line cells
197. Nectosome 0 – absent 1 – present
198. Pneumatophore 0 – absent 1 – present
199. Planula 0 – absent 1 – present
200. Planula ciliation 0 – absent 1 – present
201. Number of endodermal cells of the planula 0 – variable 1 – constant, n=16
202. Glandular cells in the planula 0 – absent 1 – present
203. Nervous cells in the planula 0 – absent 1 – present
204. Relationship between axes of planula and adult 0 – oral-aboral axis in the adult derived from the longitudinal axis of the planula 1 – oral-aboral axis in the adult derived from the transverse axis of the planula
205. Polypoid phase In previous matrix *Aegina* and *Halitrephes* had been incorrectly coded as the presence of a polypoid phase (Zhao et al., 2019), we now rescored it as absent in these two taxa and newly sampled *Halammohydra*, and accordingly change other characters contingent on the presence of a polypoid phase to inapplicable in these three taxa. 0 – absent 1 – present
206. Polyp life mode In all specimens, *Conicula* is living in solitary, neither with no signs of attaching to/with one another, or no stolon or other connected structures present. 0 – solitary 1 – colonial
207. Polymorphic polyps 0 – absent 1 – present
208. **Polyp dominant\*** Anthozoans only have a polyp phase. Although most medusozoans have polyp and medusa phases, the two phases are not equally important during the life cycles. The medusa phase is dominant in scyphozoans, cubozoans and staurozoans, while the polyp phase is generally dominant in hydrozoans. 0 – absent 1 – present
209. Stalk/peduncle in polyp This character has changed inapplicable in the bilaterians which are here regarded as no polyps present. *Conicula* is scored as present for a peduncle based on a narrower ribbon-like structure extending from the aboral end in the fossil material. 0 – absent 1 – present
210. Pedal disc 0 – absent 1 – present
211. Coelenteron Several lines of evidence indicate that *Conicula* possesses coelenteron-like features, including a blind cavity lined by mesenteries and an anthozoan-like pharynx. We score the coelenteron is present in *Conicula* 0 – absent 1 – present
212. Actinopharynx Actinopharynx refers to a short, muscular tubular passageway located between the mouth and gastric cavity in a polyp, which is formed by the invagination of the epidermis (Daly et al., 2007). *Conicula* has a distinct tubular structure similar to the actinopharynx in some aspects, including shape, topological location and dimension. We score actinopharynx as present in *Conicula*. 0 – absent 1 – present
213. Siphonoglyph This character was repeated in the previous matrix (characters 145 and 199) (Zhao et al., 2019). We retain the ‘character 199’ as it was correctly coded siphonoglyph as absent in *Corynactis*, *Montastraea* and *Porites* based on Daly et al. (Daly et al., 2003). 0 – absent 1 – present
214. Number of siphonoglyphs 0 – one 1 – more than one
215. Gastric cavity partitioned by mesentery The dark longitudinal lines preserved only in the region of gastric cavity indicate the presence of mesentery in *Conicula*. 0 – absent 1 – present
216. **Mesenterial filament \*** We split the character ‘mesenteric filament’ in the previous matrix (character 201) (Zhao et al., 2019) into two separate characters. One is for the presence of mesenterial filament and the other one is considering the number of mesenterial filament strips. Mesenterial filament refers to the thickened, cordlike margin armed with cnidae, cilia and gland cells, which occurs in the free inner edge of each mesentery below the pharynx (Brusca et al., 2016). The gastric septa of medusozoan polyp lack the mesenterial filaments. 0 – absent 1 – present
217. Number of mesenterial filament strips 0 – two strips 1 – three strips 2 – one strip
218. Ciliated tract on mesenterial filament 0 – absent 1 – present
219. Acontia Acontia are long thread-like extensions of the lower ends of mesenterial filaments, which are armed with numerous stinging cells (Lam et al., 2017). We rescore it as inapplicable in these taxa without mesenterial filaments. 0 – absent 1 – present
220. **Number of mesenteries\*** The sampled hexacorallians have more than twelve mesenteries in total except *Antipathes* which has only ten mesenteries. The sampled octocorallians have eight mesenteries, which is also scored as present in *Cambroctoconus* (Park et al., 2011). The sampled scyphozoans, cubozoans and staurozoans have four gastric septa that are also appeared in *Eoconularia* (Jerre, 1994). The estimated number of mesenteries is up to 28 in *Conicula* based on fossil material, and therefore we code *Conicula* as present for more than twelve mesenteries. 0 – more than twelve 1 – ten 2 – eight 3 – four
221. Mesentery in polyp Cubopolyps have gastrodermal folds instead of the true gastric septa (Marques and Collins, 2004), we therefore code the mesentery is absent in cubopolyps (Straehler-Pohl and Jarms, 2011). This character is rescored as inapplicable in *Aegina*, *Halitrephes* and *Halammohydra* since they develop directly to the medusae phase with no polyps present. 0 – absent 1 – present
222. **Coupled mesenteries\*** We follow Daly et al. (Daly et al., 2003) to split the character ‘Pairing of mesentery’ of the previous matrix (Zhao et al., 2019) into two separate characters (coupled mesenteries and paired mesenteries), and score them accordingly. 0 – absent 1 – present
223. **Paired mesenteries\*** 0 – absent 1 – present
224. Mesentery pair morphology 0 – members the same size 1 – members differ in size
225. Paired secondary cycle 0 – absent 1 – present
226. **Perfect mesenteries\*** This character is added primarily to establish character polarity before scoring the independent character ‘number of perfect mesenteries’. We score the perfect mesenteries as present in *Conicula*, because the presence of perfect mesenteries is presumably contingent on the presence of actinopharynx (Daly et al., 2003), a feature appeared in *Conicula*. 0 – absent 1 – present
227. **Perfect mesenteries only\*** This character is adopted from the character ‘Types of mesentery’ of the previous matrix from Zhao et al. (Zhao et al., 2019) with only considering the state ‘only perfect mesenteries’. 0 – absent 1 – present
228. Number of perfect mesenteries 0 – eight 1 – six or multiple of six
229. Directive mesentery In this matrix, we assume the gastric septa of medusozoans are homologous to mesenteries of anthozoans, and therefore rescore this character is absent in scyphozoans and staurozoans rather than inapplicable. 0 – absent 1 – present
230. Number of directive mesentery pairs 0 – one pair 1 – two pairs
231. Gonads on mesenteries of 1st cycle 0 – absent 1 – present
232. Gonads on mesenteries of 2nd and subsequent cycles 0 – absent 1 – present
233. Number of tentacles in polyp It is challenged to determine the exact number of tentacles in *Conicula*. With the incompletely exposed tentacles can be counted as up to 9, the total number of tentacles in *Conicula* is estimated to more than twenty. The Chengjiang (Ou et al., 2015) and Burgess Shale (Conway Morris and Collins, 1996) ctenophores lack evidence of tentacles, we rescore the characters that are contingent on the possession of tentacles are all inapplicable in these fossil taxa. 0 – six 1 – eight 2 – twelve 3 – sixteen 4 – eighteen 5 – more than twenty
234. Structure of polyp tentacles The polyp tentacle of *Hydra* is now coded as hollow following Ruppert et al. (Ruppert et al., 2004). It is unknown the structure of polyp tentacles in *Conicula* and other fossil groups. 0 – hollow 1 – solid
235. Arrangement of tentacles 0 – scattered 1 – one cycle 2 – more than one cycle
236. Two-tentacle polyp stage 0 – absent 1 – present
237. Tentacles retractile 0 – non retractile 1 – retractile
238. Tentacle/coelenteron relationship 0 – one tentacle per endocoel and per exocoel 1 – one tentacle per exocoel, multiple per endocoel
239. Catch tentacles Catch tentacles is a special type of tentacle longer than ordinary tentacles, probably using for social behaviour, which is only present in *Metridium* (Williams, 1975) of our sampled taxa. We score *Conicula* and dinomischids (Zhao et al., 2019) as absent for catch tentacles based on its gross appearance of tentacles. 0 – absent 1 – present
240. Acrospheres 0 – absent 1 – present
241. Marginal spherules 0 – absent 1 – holotrichous
242. Acrorhagi 0 – absent 1 – present
243. Organisation of the nervous system 0 – nets 1 – with nerve rings
244. Canal system in polyp 0 – absent 1 – present
245. Gastrodermic musculature 0 – not in bunches 1 – organised in bunches of gastrodermal origin 2 – organised in bunches of ectodermal origin
246. Mesogleal sphincter 0 – absent 1 – present
247. Ectodermal longitudinal muscle location 0 – tentacles and oral disc only 1 –whole body
248. Basilar musculature 0 – absent 1 – present
249. Retractor muscle 0 – weak 1 – defined
250. Parietal muscle 0 – absent 1 – present
251. Mesogleal lacunae 0 – absent 1 – present
252. Actinula This is an easily confused term as it widely uses to describe the free-moving larva stage of Trachymedusae and Anthomedusae, but the two are not homologous (Bouillon and Boero, 2000; Petersen, 1990). Here, we refer the character actinula only to the larva stage of Anthomedusae, and accordingly score it present in *Candelabrum* and *Ectopleura* (Petersen, 1990). 0 – absent 1 – present
253. Ephyrae The ephyra is a distinguishing characteristic of living scyphozoans and is absent in staurozoans and cubozoans. Based on fossil material, *Olivooides* may also have an ephyra stage during the development process (Dong et al., 2013). 0 – absent 1 – present
254. **Marginal lappet type\*** The marginal lappet of ephyrae generally comprises two portions (lappet stem and rhopalial lappet) as seen in *Aurelia* and *Rhizostoma*, while the lappet stem is absent in *Linuche*, *Atorella* and *Nausithoe* (Straehler-Pohl and Jarms, 2010). It is unknown the type of marginal lappet of *Olivooides* based on the fossil material. 0 – without lappet stem 1 – with lappet stem
255. **Number of marginal lappets\*** *Aurelia* and *Rhizostoma* have eight marginal lappets. Coronate *Linuche* and *Nausithoe* have sixteen lappets whereas *Atorella* possesses twelve lappets (Daly et al., 2007). *Olivooides* probably has five lappets (Dong et al., 2013). 0 – five 1 – eight 2 – twelve 3 – sixteen
256. Medusoid phase The medusoid phase is one of the biphasic life cycles of cnidarians, referring to the presence of free-living medusa. It is uncertain in *Conicula* as all specimens are in the polypoid phase and the medusoid form is still yet to be determined. Considering the presence of ephyra (Dong et al., 2013), it is therefore scored this character as present in *Olivooides*. 0 – absent 1 – present
257. Location of medusa formation Because of the presence of ephyra (Dong et al., 2013), we score *Olivooide*s as producing medusa from apical or oral location. 0 – lateral budding from an entocodon 1 – apical or oral 2 – direct development without polyp stage
258. Type of apical medusa formation Unlike the traditional opinion, several cubozoans also have strobilation-like metamorphosis process, such as the newly sampled taxon *Carukia* (Courtney et al., 2016). The ephyra is one of the results of strobilation, we score *Olivooides* as strobilation because of the possible fossil record of ephyra (Dong et al., 2013). 0 – strobilation 1 – without transverse fission
259. Strobilation type Cubozoan *Carukia* produces monodisc strobilation (Courtney et al., 2016). *Aurelia*, *Linuche* and *Nausithoe* have polydisc strobilation (Helm, 2018), while *Rhizostoma* has both polydisc and monodisc strobilation (Holst et al., 2007). *Olivooides* has two tightly fused ephyrae (Dong et al., 2013), probably indicating the presence of polydisc strobilation. 0 – polydisc 1 – monodisc
260. Adult medusoid shape Although the state ‘3 – actinuloid’ existed in the previous matrix, none of taxa has been scored as ‘actinuloid’ (Zhao et al., 2019). Here a new taxon *Halammohydra* is added into the sampled taxa, and is scored as the presence of actinuloid (Clausen, 1967). 0 – bell 1 – pyramidal 2 – cubic 3 – actinuloid
261. Shape of horizontal cross-section of the medusa 0 – circular 1 – four-part symmetry
262. Development of the umbrella The sampled taxa in previous matrix (Zhao et al., 2019) were all coded as present for a fully developed umbrella except the score of inapplicable and unknown. Here we add a new taxon *Halammohydra*, of which the medusa develops only the aboral cone (Clausen, 1967). 0 – fully developed 1 – aboral cone
263. **Umbrellar size\*** Scyphomedusa is named as the ‘true jellyfishes’ that have a well-developed umbrella up to 2 metres (e.g. *Cyanea*), while hydromedusa is mostly small. Cubomedusa and stauromedusa are also small compared with scyphomedusa, but is larger than hydromedusa (Brusca et al., 2016). 0 – mostly small (up to 10cm in diameter) 1 – small (up to 30cm in diameter) 2 – big (up to 2m in diameter)
264. Umbrellar margin 0 – smooth and continuous 0 – lobed
265. Rhopalia/rhopalioids 0 – absent 1 – present
266. Complex eyes in rhopalia 0 – absent 1 – present
267. Statocysts 0 – absent 1 – present
268. Statocyst origin 0 – endodermic 1 – ectodermic
269. Statolith composition 0 – MgCaPO4 1 – CaSO4
270. Ocelli in medusa 0 – absent 1 – present
271. Giant fibre nerve net (GFNN) in medusae The character was duplicated in the previous matrix (characters 23, 147 and 258) (Zhao et al., 2019), and here we score it inapplicable in *Hydra* and *Candelabrum* because of the absence of the medusoid phase. 0 – absent 1 – present
272. Nerve ring in medusa 0 – absent 1 – present
273. Number of rings in nerve ring 0 – one 1 – two
274. Manubrium 0 – absent 1 – present
275. Gastric filaments 0 – absent 1 – present
276. **Gastric saccule\*** This new character is widely present in the chirodropids of cubozoans (Daly et al., 2007). It is coded as absent in *Carybdea*, *Carukia* and other medusozoans. 0 – absent 1 – present
277. Coronal muscle 0 – well-developed 1 – marginal and tiny
278. **Longitudinal muscles in the peduncle\*** This character is contingent on the presence of peduncle, a character generally present in staurozoans. *Haliclystus* has longitudinal muscles in the peduncle but it is absent in *Craterolophus* (Miranda et al., 2016). 0 – absent 1 – present
279. Pedalium of coronate type 0 – absent 1 – present
280. Pedalium of cubozoan type 0 – absent 1 – present
281. **Pedalial branching\*** The branched pedalia bearing numerous tentacles appear in the chirodropids of cubozoans (Daly et al., 2007). This new character is present in *Chiropsalmus* (Gershwin, 2006) of our sampled taxa. 0 – absent 1 – present
282. Velum We rescore the velum as absent in *Obelia* (Bouillon and Boero, 2000). 0 – absent 1 – present
283. **Claustrum\*** The claustrum refers to a membrane constituted by layers of mesoglea and gastrodermis that divides the gastric cavity, which is exclusive to some of the staurozoans and is absent in other cnidarians, including the cubozoans (Miranda et al., 2017). In our sampled taxa, claustrum is present in *Craterolophus* and is absent in *Haliclystus* (Miranda et al., 2016). 0 – absent 1 – present
284. Tentacles in medusa 0 – absent 1 – present
285. Structure of medusa tentacle *Obelia* is rescored as the presence of solid marginal tentacles (Bouillon and Boero, 2000). 0 – hollow 1 – solid
286. Shape of medusa tentacle 0 – filiform 1 – capitate
287. Tentacular bulbs Following the recent studies (Holst et al., 2021; Miranda et al., 2013; Miranda et al., 2016), we now score the tentacular bulbs as absent in staurozoans. 0 – absent 1 – present
288. Tentacular insertion 0 – at umbrellar margin 1 – away from margin
289. Number of tentacular whorls In the matrix of Zhao et al. (Zhao et al., 2019), most medusozoans were coded as one tentacular whorl, but none of the taxa was scored as present for two tentacular whorls. We now add a new taxon *Halammohydra*, which has two whorls of tentacles (Clausen, 1967). 0 – one whorl 1 – two whorls
290. Oral arms with suctorial mouths 0 – absent 1 – present
291. Mesentery in medusa 0 – absent 1 – present
292. Mesenteric shape in medusa 0 – straight 1 – Y-shaped
293. Perradial mesenteries 0 – absent 1 – present
294. Radial canals 0 – absent 1 – present
295. Circular canal 0 – absent 1 – present
296. Circular canal partial 0 – absent 1 – present
297. Peripheral canal system 0 – absent 1 – present
298. Velarium 0 – absent 1 – present
299. Velar canals 0 – absent 1 – present
300. Frenulae 0 – absent 1 – present
301. Coronal furrow 0 – absent 1 – present
302. Gonadal location Unlike the score in the Zhao et al. (Zhao et al., 2019), we add a new state (2 – both sides of gastric septa) to define the gonadal location of staurozoans (Holst et al., 2021; Miranda et al., 2013), since the radial canals are absent in staurozoans. 0 – manubrium 1 – radial canals 2 – both sides of gastric septa
303. Urticant rings 0 – absent 1 – present
304. Peronia 0 – absent 1 – present

## Notes

### Competing Interest Statement

The authors have declared no competing interest.

